# Chemoresistance of *TP53* mutant AML requires the mevalonate byproduct, GGPP, for regulation of ROS and induction of a mitochondria stress response

**DOI:** 10.1101/2024.06.07.597976

**Authors:** Sarah J. Skuli, A’Ishah Bakayoko, Marisa Kruidenier, Bryan Manning, Paige Pammer, Akmal Salimov, Owen Riley, Gisela Brake-Sillá, Michael Bowman, Leslie N. Martinez-Gutierrez, Roberta Buono, Madhuri Paul, Estelle Saland, Sarah Wong, Jimmy Xu, Eva Nee, Ryan Hausler, Colin Anderson, Julie A. Reisz, Angelo D’Alessandro, Catherine Lai, Kara N. Maxwell, Jean-Emmanuel Sarry, David A. Fruman, Clementina Mesaros, Brian Keith, M. Celeste Simon, Pamela J. Sung, Gerald Wertheim, Nicolas Skuli, Robert L. Bowman, Andrew Matthews, Martin Carroll

## Abstract

Acute myeloid leukemia (AML) with mutations in the tumor suppressor gene, *TP53* (*TP53*^mut^ AML), is fatal with a median survival of only 6 months. RNA sequencing on purified AML patient samples show *TP53*^mut^ AML has higher expression of mevalonate pathway genes. We retrospectively identified a survival benefit in *TP53*^mut^ AML patients who received chemotherapy concurrently with a statin, which inhibits the mevalonate pathway. Mechanistically, *TP53*^mut^ AML resistance to standard AML chemotherapy, cytarabine (AraC), correlates with increased mevalonate pathway activity and a mitochondria stress response with increased mitochondria mass and oxidative phosphorylation. Pretreatment with a statin reverses these effects and chemosensitizes *TP53*^mut^ AML cell lines and primary samples *in vitro* and *in vivo*. Mitochondria-dependent chemoresistance requires the geranylgeranyl pyrophosphate (GGPP) branch of the mevalonate pathway and novel GGPP-dependent synthesis of glutathione to manage AraC-induced reactive oxygen species (ROS). Overall, we show that the mevalonate pathway is a novel therapeutic target in *TP53*^mut^ AML.

**Significance:** Chemotherapy-persisting *TP53*^mut^ AML cells induce a mitochondria stress response that requires mevalonate byproduct, GGPP, through its novel role in glutathione synthesis and regulation of mitochondria metabolism. We provide insight into prior failures of the statin family of mevalonate pathway inhibitors in AML. We identify clinical settings and strategies to successfully target the mevalonate pathway, particularly to address the unmet need of *TP53*^mut^ AML.

## Introduction

In the field of leukemia, *TP53* mutant acute myeloid leukemia (*TP53*^mut^ AML) is the paramount clinical challenge. In *TP53*^mut^ patients, standard AML therapies, including cytarabine (AraC)-based regimens as well as hypomethylating (HMA) agents with or without the Bcl2 inhibitor, venetoclax (Ven), all lead to a similar and dismal median overall survival of only 6-9 months(1,2). Thus, international guidelines recommend a clinical trial as the first approach to treatment of *TP53*^mut^ AML. Recent studies suggest a benefit of allogeneic hematopoietic stem cell transplant if patients can achieve an initial complete response(1,3). However, the vast majority of patients with *TP53*^mut^ AML are either primary refractory or relapse quickly following an initial response to therapy. Furthermore, relatively few patients achieve a minimal residual disease-negative remission as defined by the persistence of a *TP53* mutation by next generation sequencing(4). Thus, the primary etiology of poor outcomes of *TP53*^mut^ AML is chemotherapy resistance. We propose that the potential for chemoresistance is not binary but exists on a spectrum across AML. *TP53*^mut^ AML is the ultimate model in which mechanisms of chemoresistance will be most readily identified. Importantly, novel strategies that reverse chemoresistance in *TP53*^mut^ AML may be applicable to other chemoresistant AMLs.

Our group and others have demonstrated that AML requires enhanced mitochondria oxidative phosphorylation (OXPHOS) to survive chemotherapy, including AraC(5–7). It has also been proposed that inhibition of OXPHOS is the primary driver of synergy in Ven/HMA combinations, with increased mitochondria fatty acid oxidation being a mechanism of therapy resistance(8,9). The relationship between chemoresistance and increased mitochondria activity is especially interesting in the context of *TP53*^mut^ AML, since loss of p53 leads to increased glycolysis, OXPHOS, and mitochondria biogenesis in both solid tumors and a recent AML model(10–12). However, the direct relationship between increased mitochondria activity and resistance to chemotherapy in *TP53*^mut^ AML has yet to be addressed, as many studies on AML therapy response have been performed without regard to *TP53* mutation status. Moreover, direct targeting of mitochondria metabolism to reverse chemotherapy resistance has been limited by toxicity due to essential mitochondria functions in non-malignant cells(13). Novel strategies are needed to identify and target pathways that contribute to mitochondria metabolism primarily and specifically in cancer cells.

Here, we demonstrate that transcriptional analysis of primary human AML samples shows the mevalonate pathway is overexpressed in *TP53*^mut^ compared to more chemosensitive *TP53* wildtype (*TP53*^WT^) samples. Maintaining cholesterol homeostasis is a critical cellular function as recently reviewed(14,15). While cholesterol can be imported into the cell, it can also be synthesized *de novo* by the mevalonate pathway, a complex and tightly regulated metabolic pathway. The mevalonate pathway branches at the level of farnesyl pyrophosphate (FPP), forming either isoprenoids, such as geranylgeranyl pyrophosphate (GGPP), or other cholesterol derivatives. Importantly, there is significant cross-talk and interdependence of mitochondria metabolism with the mevalonate pathway(16–19). Furthermore, the mevalonate pathway is necessary for tumorigenesis and metastatic potential of multiple *TP53*^mut^ solid tumor models(20–24). In AML, several studies have implicated the mevalonate pathway in chemoresistance(25–29). Early phase clinical trials investigating the addition of statins to AraC-based induction therapy showed the most promising benefit in patients with relapsed/refractory, poor risk AML(30–32) but did not lead to statin incorporation into AML standard therapy. Notably, *TP53*^mut^ AML patients should be enriched in this relapsed/refractory AML subset, but sequencing of *TP53* in AML was not routinely performed at the time of the study. Altogether, the mechanism of action and the role of *TP53* in metabolic regulation in AML remain poorly understood.

We hypothesized that the mevalonate pathway is required for mitochondria-dependent chemoresistance in *TP53*^mut^ AML. We used multiple, FDA-approved and commonly prescribed mevalonate pathway inhibitors, particularly the statin drug class, as a tool to determine the role of the mevalonate pathway in the chemoresistance of *TP53*^mut^ AML. Furthermore, we addressed this hypothesis with multiple orthogonal approaches, including previously unpublished, multi-institutional, retrospective clinical data, novel, isogenic *TP53*^mut^ AML cell lines and primary *TP53*^mut^ AML patient samples. Our studies indicate that the mevalonate pathway should be targeted in chemotherapy resistant *TP53*^mut^ AML and identifies novel therapeutic strategies to achieve chemosensitization.

## Results

### Primary *TP53*^mut^ AML RNA sequencing shows upregulation of a cholesterol synthesis gene signature

To elucidate biological differences between *TP53*^mut^ and *TP53*^WT^ AML, we performed RNA sequencing on a pure population of untreated, primary AML blasts obtained by flow sorting viably frozen mononuclear cells for a human CD45 and CD33 positive immunophenotype. Primary AML samples were selected by *TP53* mutation status and overall survival with full clinical annotation is summarized in Supplemental Table 1A. 9 patients with *TP53*^mut^ AML were selected, of which 8 had at least 1 missense mutation in *TP53* with the remaining patient having a *TP53* frameshift mutation, all with variant allele fractions (VAF) greater than 40% (Supplemental Table 1A). The median overall survival of the *TP53*^mut^ AML patients was 65 days (Supplemental Fig1A), consistent with the literature. An additional 21 *TP53*^WT^ AML patient samples with either a “good” overall survival defined as greater than 5 years (n=9; median overall survival not reached) or a “bad” overall survival of less than 2 years (n=12, median overall survival 294 days) were also included (Supplemental Fig 1A). Principle component analysis comparing *TP53*^mut^ *versus TP53*^WT^ good outcome *versus TP53*^WT^ bad outcome AML suggested the *TP53*^mut^ AML patient samples primarily clustered together whereas the *TP53*^WT^ AML patient samples were more heterogeneous (Fig 1A). For subsequent studies, all *TP53*^WT^ AML patient samples were grouped together to compare *TP53*^mut^ vs *TP53*^WT^ AML [Differential gene expression and gene set enrichment analysis (GSEA) summarized in Supplemental Table 1B-C]. Single sample GSEA (ssGSEA) demonstrated primary *TP53*^mut^ compared to *TP53*^WT^ AML had higher expression of the LSC17 signature(33) but lower expression of *TP53* targets(34) (Fig 1B-C). Notably, multiple gene sets involved in cholesterol synthesis were upregulated in *TP53*^mut^AML (Fig 1B-C; Supplemental Table 1D). These signatures were recapitulated in a cohort of primary *TP53*^mut^ (n=17) *versus TP53*^WT^ (n=87) AML samples identified in the BeatAML dataset, with a key difference being RNA sequencing was performed on ficoll-purified mononuclear cell fractions without cell sorting for a pure blast population (Supplemental Fig 1C-D). Of note, the *TP53*^mut^ AML patients in the BeatAML subset had an equally poor overall survival of 165 days, which was significantly lower than the median overall survival of 520 days in the *TP53*^WT^ cohort (p < 0.0001; Supplemental Fig 1B).

**Figure 1:**
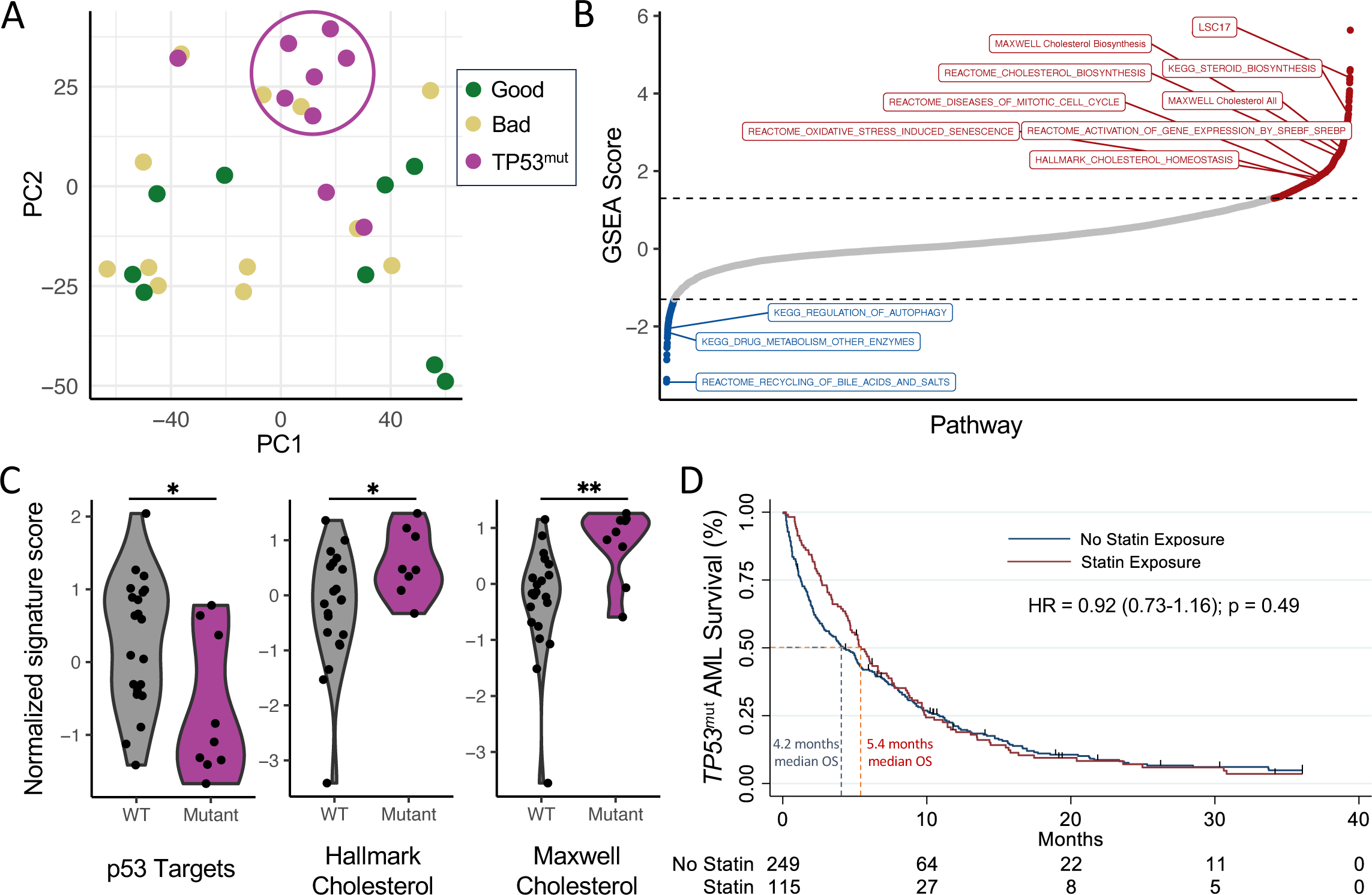
Untreated, primary *TP53*^mut^ AML are enriched for cholesterol synthesis gene expression, and show improved overall survival when statins are co-administered with AML therapy. RNA sequencing was performed on highly purified, flow-sorted blasts of primary human AML diagnostic samples from patients with *TP53*^mut^ (n=9), *TP53*^WT^ “bad” outcome AML (n=12), and *TP53*^WT^ “good” outcome AML (n=9) with subsequent analysis of the (A) principle component analysis, (B) GSEA scores from the Hallmark and other denoted gene sets, and (C) single sample GSEA with calculation of normalized signature scores for “TP53 Targets,” “Hallmark Cholesterol,” and “Maxwell Cholesterol.” Statistical analysis by Student’s T Test. (D) Kaplan-Meier survival estimates from a retrospective study of newly diagnosed *TP53*^mut^ AML patients (n=364) who did or did not receive a concurrent statin during induction therapy (n=115 and n=249, respectively).

### Statins improve overall survival of *TP53*^mut^ AML patients undergoing induction therapy

Statins are a class of drugs that inhibit the mevalonate pathway and are one of the most commonly prescribed drugs for cardiovascular disease. Thus, a retrospective analysis was performed to determine the impact of concomitant statin with induction therapy on overall survival in patients with newly diagnosed *TP53*^mut^ AML. Retrospective chart reviews were performed on 364 *TP53*^mut^ AML patients treated at one of two tertiary centers on the East Coast between 2013-2023 and demographic and clinical data was recorded (Supplemental Table 1E). 32% of patients received a statin during induction therapy for *TP53*^mut^ AML. Statin-treated patients were older (age 70 vs 68, p = 0.004) and had more indications for a statin, such as a history of coronary artery disease and stroke (Fisher’s exact < 0.001), but were otherwise well matched. An unadjusted Kaplan-Meier survival estimate suggested a non-significant 1.2-month improvement in overall survival in patients who received a statin with induction therapy (Fig 1D). Statins appear to offer a significant benefit initially, but the survival curves overlap again by 6 months. These data provided a rationale to investigate the role of the mevalonate pathway in *TP53*^mut^ AML chemotherapy resistance using statins as a tool to manipulate the mevalonate pathway.

### *TP53*^mut^ AML cell lines require the mevalonate pathway to survive AML chemotherapy, AraC

Isogenic *TP53*^mut^ AML cell lines with complete loss (M14-Mut1, M14-Mut3) or missense-like mutation (M14-Mut2, M14-Mut4) of the p53 protein were developed using CRISPR/Cas9 editing of the AML cell line, MOLM-14, with published guides(35,36) and single cell cloning *via* serial dilution. Full molecular annotation of the parental and isogenic cell lines is available in Supplemental Table 1F. To determine the impact of loss or mutation-induced dysfunction of p53 on chemosensitivity, we treated the isogenic *TP53*^mut^ (M14-Mut1, M14-Mut2, M14-Mut3, M14-Mut4) and *TP53*^WT^ (M14-WT1) AML clones with increasing concentrations of AraC for 24 hours and assessed cell viability by flow cytometry after staining for AnnexinV and 7AAD. The *TP53*^mut^ AML clones had consistently higher viability than the *TP53*^WT^ AML clone, M14-WT1, at the same AraC concentrations indicating that the *TP53*^mut^ AML clones are more chemoresistant and an appropriate *TP53*^mut^ AML model (Fig 2A). We then validated that only the M14-WT1 clone exhibits a rapid and significant p53-dependent increase in downstream p53 target, p21, following AraC treatment (Supplemental Fig 2A). To determine the effect of inhibition of the mevalonate pathway, we pretreated the isogenic *TP53*^mut^ and *TP53*^WT^ AML clones with the mevalonate pathway inhibitor, rosuvastatin, for 24 hours prior to adding AraC for an additional 24 hours and assessed viability by flow cytometry. Importantly, rosuvastatin chemosensitized the *TP53*^mut^ AML clones to AraC (Fig 2B). Subsequent XTT viability assays showed that the combination of AraC and rosuvastatin was synergistic in *TP53*^mut^ (M14-Mut1 mean bliss score 30) and *TP53*^WT^ (M14-WT1 mean bliss score 36) AML clones using the SynergyFinder web-based tool, with a score greater than 10 being likely synergistic, −10 to 10 likely additive, and less than −10 likely antagonistic(37) (Supplemental Fig 2B). A second statin, pitavastatin, also chemosensitized *TP53*^mut^ AML clones to AraC (Supplemental Fig 2C) indicating that statin chemosensitization is a class-effect.

**Figure 2:**
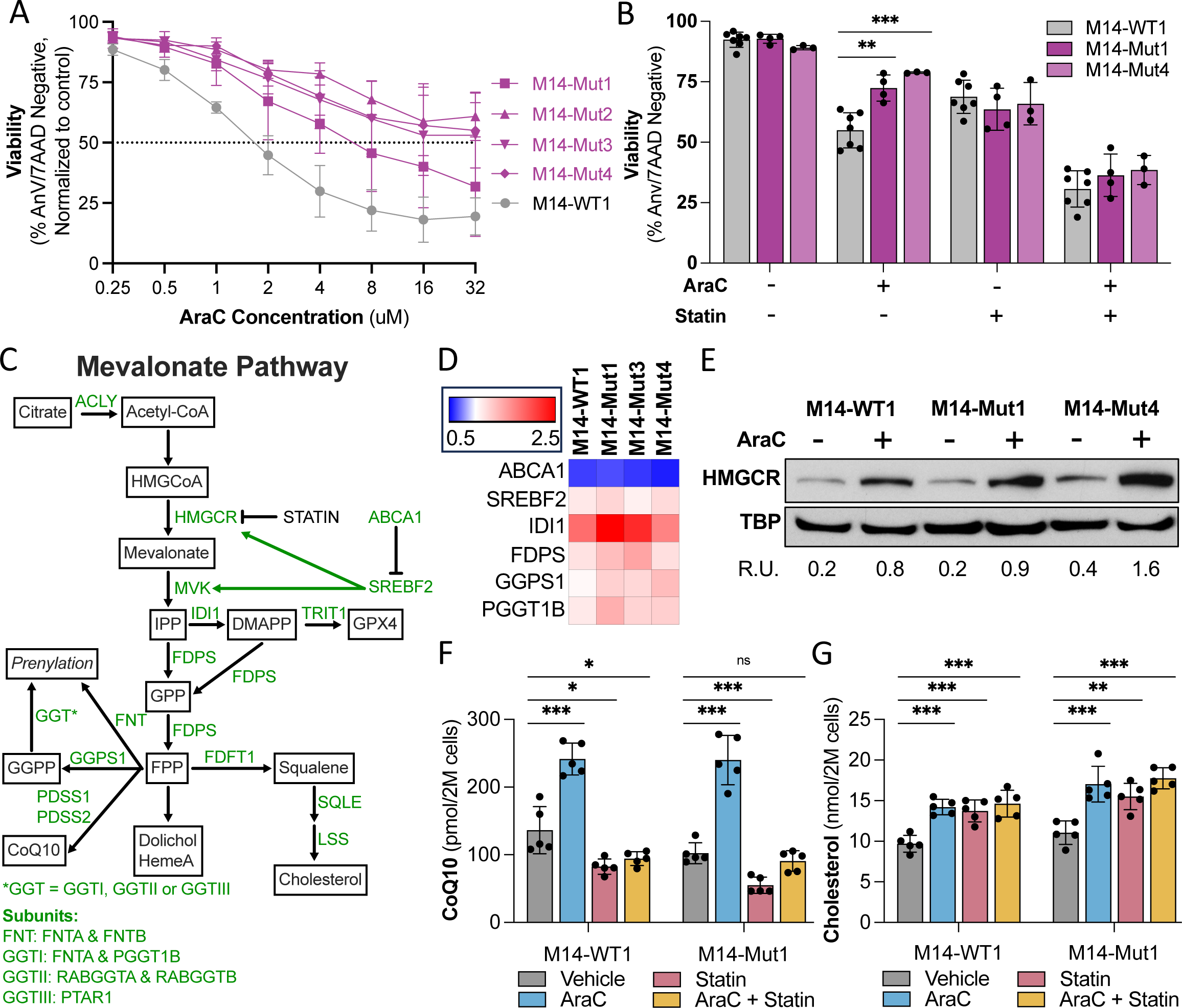
Chemoresistant *TP53*^mut^ AML cell lines upregulate the mevalonate pathway in response to AraC. (A) Cell viability of isogenic *TP53*^mut^ AML clones treated with increasing concentrations of AraC for 24 hours and assessed by flow cytometry following staining with AnnexinV and 7AAD (n=3). (B) Cell viability of representative isogenic *TP53*^mut^ and *TP53*^WT^ AML clones pretreated with 24 hours of rosuvastatin (50uM) followed by an additional 24 hours of AraC (1uM) and assessed by flow cytometry following staining with AnnexinV and 7AAD (n=7). (C) Schema of the mevalonate pathway with key enzymes highlighted in green and byproducts in squares. (D) Gene expression of select mevalonate pathway genes in isogenic *TP53*^mut^ and *TP53*^WT^ AML clones following 24 hours of AraC (1uM) treatment, normalized to GAPDH, and compared to the vehicle for each clone as measured by qRT-PCR (n=3). (E) Protein expression of HMGCR normalized to total binding protein (TBP) in isogenic *TP53*^mut^ and *TP53*^WT^ AML clones treated with vehicle or 18 hours of AraC (1uM), performed by western blot and quantified by imageJ as presented in relative units (R.U.) of HMGCR to TBP. (F) CoQ10 as pmol per 2 million cells and (G) cholesterol as nmol per 2 million cells in the M14-WT1 and M14-Mut1 clones in the same conditions as (B) and quantified by LC-HRMS (n=5). Statistical analysis by Student’s T Test. p-values: * = <0.05, ** = <0.01, *** = <0.001. n is the number of replicates.

### Chemoresistant *TP53*^mut^ AML cell lines upregulate the mevalonate pathway in response to AraC

Representative isogenic *TP53*^mut^ and *TP53*^WT^ AML clones were used to assess the impact of p53 loss on the mevalonate pathway at baseline and in response to AraC, with key mevalonate pathway byproducts and genes highlighted in the schema in Figure 2C. Mevalonate pathway gene expression was first assessed by qRT-PCR. AraC induces time-dependent upregulation of multiple genes in the mevalonate pathway as well as the pathway’s primary transcription factor, SREBF2, in both *TP53*^mut^ and *TP53*^WT^ AML clones (Fig 2D, Supplemental Fig 2D for full timecourse). HMGCR and SREBF2 are also regulated at the post-translational level(38). HMGCR protein expression, as determined by western blot, is significantly increased by AraC in all clones tested and highest in the *TP53*^mut^ AML clones (Fig 2E, Supplemental Fig 2E). While the immature or inactive endoplasmic-reticulum bound form of SREBF2 is mildly increased in response to AraC, there is no detectable increase in the mature or active form of SREBF2 in whole cell lysates (Supplemental Fig 2F). Finally, mevalonate pathway byproducts, CoQ10 and free cholesterol, were assessed by liquid chromatography, high-resolution mass spectrometry (LC-HRMS). A significant increase in CoQ10 and cholesterol was detected in AraC-treated isogenic *TP53*^mut^ and *TP53*^WT^ AML clones (Fig 2F-G, respectively). Rosuvastatin pretreatment abrogated the increase in CoQ10 (Fig 2G), while rosuvastatin pretreatment alone or in combination with AraC was associated with a significant increase in cholesterol (Fig 2H). Altogether, these data suggest that the isogenic clones primarily synthesize the isoprenoid CoQ10 tail *via* the mevalonate pathway, but respond to mevalonate pathway inhibition with increased cholesterol uptake. Notably, key differences in the isogenic *TP53*^mut^ vs *TP53*^WT^ AML clones in regards to levels of mevalonate gene, protein and byproduct expression are most pronounced following stress induction such as that induced by AraC.

### Mevalonate pathway-driven OXPHOS and regulation of ROS correlate with AraC resistance in *TP53*^mut^ AML cell lines

Our group and others have demonstrated that AML AraC resistance correlates with increased mitochondria OXPHOS. While p53 is known to regulate metabolism, the impact of *TP53* mutations on mitochondria metabolism at baseline and in response to AraC in AML is not well understood. Thus, we aimed to assess mitochondria metabolism in our isogenic *TP53*^mut^ *versus TP53*^WT^ AML clones. In response to AraC, only *TP53*^mut^ AML clones induce a significant increase in basal and maximum uncoupler-induced oxygen consumption [Basal OXPHOS: Fold-change of 1.2 (p = 0.05) for M14-WT1 and 1.5 (p < 0.001), 1.9 (p < 0.001), 2.1 (p = 0.002), and 1.9 (p = 0.03) for M14-Mut1, M14-Mut2, M14-Mut3, and M14-Mut4, respectively; Maximum-uncoupler induced OXPHOS: Fold-change of 1.3 (p = 0.04) for M14-WT1 and 2 (p < 0.001), 2 (p = 0.03), 2.7 (p < 0.001), and 2.2 (p = 0.04) for M14-Mut1, M14-Mut2, M14-Mut3, and M14-Mut4, respectively] as assessed by Seahorse technology (Fig 3A-B, Supplemental Fig 3A-B). The increase in OXPHOS is associated with an increase in mitochondria mass as determined by measuring the ratio of mitochondrial to nuclear DNA by PCR (Fig 3C). As expected, treatment of M14-WT1 with AraC was associated with a significant increase in mitochondrial reactive oxygen species (ROS) (6-fold increase compared to vehicle, p < 0.001) (Fig 3D). Correlating with chemoresistance, M14-Mut1 and M14-Mut4 exhibited a significantly lower induction of mitochondrial ROS in response to AraC (2-fold increases compared to vehicle, p < 0.001 for both) (Fig 3D). Thus, there is a paradoxical improvement in ROS management in *TP53*^MT^ AML clones despite, and potentially allowing, a significant increase in OXPHOS in response to AraC and as compared to M14-WT1 (Fig 3A-B, D). Furthermore, AraC-induced ROS and apoptosis is at least partially rescued by pretreatment with the antioxidant, glutathione (Fig 3D-E), suggesting AraC-induced ROS directly correlates to induction of cell death.

**Figure 3:**
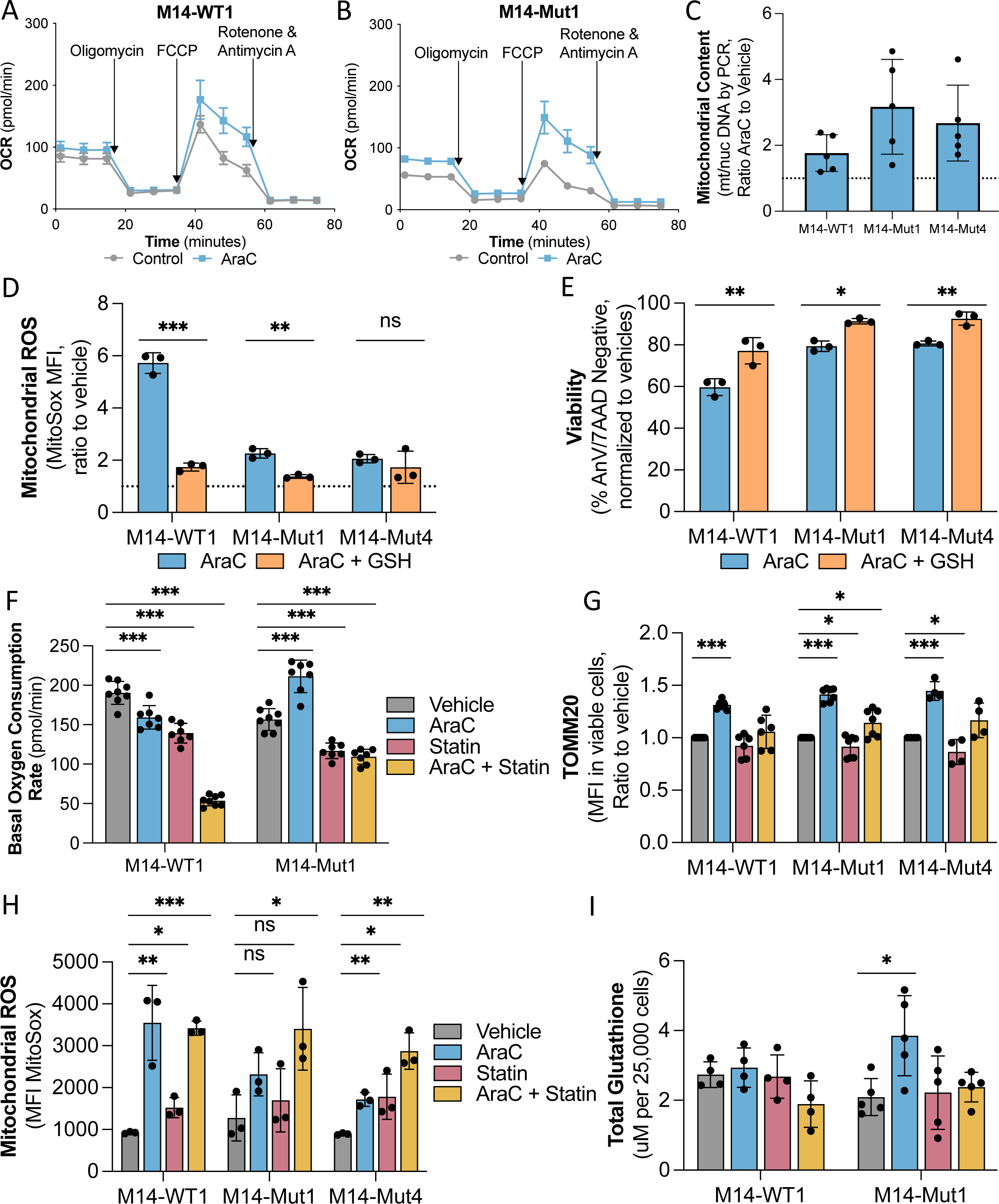
Mevalonate pathway-driven OXPHOS and regulation of ROS correlate with AraC resistance specifically in *TP53*^mut^ AML cell lines. Oxygen consumption in pmol per minute in (A) M14-WT1 and (B) M14-Mut1 treated with vehicle or AraC (1uM) for 24 hours and measured by Seahorse technology (n=7 and n=8, respectively). (C) Mitochondrial content presented as ratio of AraC to vehicle of mitochondrial to nuclear DNA measured by PCR in representative isogenic *TP53*^mut^ and *TP53*^WT^ AML clones treated with vehicle or AraC (1uM) for 24 hours. (D) Mitochondrial ROS presented as the ratio to vehicle of mean fluorescence intensity (MFI) of MitoSox in viable cells [determined by Fixable Viability Stain Red (FVS-R)] as measured by flow cytometry in representative isogenic *TP53*^mut^ and *TP53*^WT^ AML clones treated with AraC (1uM, 24 hours), glutathione (10mM, 36 hours), the combination, or their respective vehicles (n=3). (E) Viability of cells treated as per (D) and measured by flow cytometry after staining with AnnexinV and 7AAD. For F-I, representative isogenic *TP53*^mut^ and *TP53*^WT^ AML clones were treated with rosuvastatin (50uM) for a total of 48 hours with AraC (1uM) added for the last 24 hours and the following experiments subsequently performed: (F) Basal oxygen consumption rate in pmol per minute assessed by Seahorse technology (n=7), (G) TOMM20 presented as MFI in viable cells (by FVS-R) assessed by flow cytometry (n=6, n=7, n=4 for respective cell lines), (H) mitochondrial ROS presented as the ratio to vehicle of MFI of MitoSox in viable cells (by FVS-R) as measured by flow cytometry (n=3), and (I) total glutathione presented as uM per 25,000 cells (n=4, n=5 for respective cell lines). Statistical analysis by Student’s T Test. p-values: * = <0.05, ** = <0.01, *** = <0.001. n is the number of replicates.

Given multiple mevalonate pathway byproducts are particularly important for mitochondria metabolism and redox management, we next assessed the impact of pretreatment with rosuvastatin on mitochondria parameters. Rosuvastatin alone decreases basal and max OXPHOS (Fig 3F, Supplemental Fig 3C-D) and decreases TOMM20, a mitochondria-specific protein that functions as a surrogate for mitochondria mass (Fig 3G). Furthermore, inhibition of the mevalonate pathway with rosuvastatin alone increases mitochondrial ROS in all clones (Fig 3H). Importantly, in M14-Mut1 and M14-Mut4, rosuvastatin pretreatment completely abrogates the AraC-induced increase in OXPHOS and TOMM20 while increasing mitochondrial ROS to the same degree as M14-WT1 (Fig 3F-H, Supplemental Fig 3C-D). Finally, in correlation with improved AraC-induced ROS management in *TP53*^MT^ AML clones, M14-Mut1, but not M14-WT1, responds to AraC with a significant increase in total glutathione, including a specific increase in the reduced form of glutathione (Fig 3I, Supplemental Fig 3E). The increase in glutathione in M14-Mut1 in response to AraC is mevalonate pathway-dependent, as pretreatment with rosuvastatin abrogates the increase in reduced and total glutathione (Fig 3I, Supplemental Fig 3E). Altogether, this data suggests that a mitochondria stress response is key for AraC chemoresistance in *TP53*^mut^ AML and requires mevalonate pathway-dependent ROS management. Furthermore, we identify a previously undescribed role of the GGPP branch of the mevalonate pathway in glutathione synthesis.

### PDX modelling of *TP53*^mut^ AML reveals modest statin-mediated AraC chemosensitization

We established a PDX mouse model of *TP53*^mut^ AML in busulfan-conditioned NOD scid gc^-/-^ (NSG) mice. The primary sample used for the study was obtained from an untreated patient with *de novo* AML with a frameshift *TP53* mutation identified on next generation sequencing with a variant allele frequency of 90% and a complex monosomal karyotype. Following engraftment, mice were treated with vehicle, AraC (50mg/kg/day intraperitoneally on days 1-5), rosuvastatin (1mg/kg/day by oral gavage on days 1-7, the maximum dose tolerated in patients(39)) or the combination, as described in our schema (Fig 4A). Rosuvastatin was well tolerated with no detrimental effects on body weight (data not shown). Mouse bone marrow (BM) and spleen were harvested on days 8-9 and leukemic burden was determined by assessing the percentage of human CD45+/CD33+ cells by flow cytometry and quantifying with counting beads. Only the combination of AraC and rosuvastatin significantly decreased leukemic burden (Fig 4B). Consistent with *in vitro* data, surviving leukemic cells from AraC-treated mice had a significant increase in OXPHOS (Fig 4C, Supplemental Fig 4A for all seahorse tracings) and mevalonate pathway byproducts, CoQ10 (Fig 4D) and cholesterol (Fig 4E), assessed by LC-HRMS. However, rosuvastatin did not fully reverse AraC-induced OXPHOS or mevalonate pathway byproduct accumulation, suggesting inadequate inhibition of the mevalonate pathway by rosuvastatin *in vivo* (Fig 4C-E). The experiment was also performed with a different primary sample, SCXC-7575, that was obtained from a patient with relapsed, secondary AML who had previously received HMA/venetoclax and was noted to have mutations in *TP53* (R267W missense mutation, VAF 99%), *IDH2* (VAF 49%), *ASXL1* (VAF 45%), *KRAS* (VAF 47%) and *CEBPA* (49%), as well as a complex karyotype. AraC significantly reduced leukemic burden in this *TP53*^mut^ AML PDX mouse model with no additional benefit from rosuvastatin (Fig 4F). Furthermore, AraC-persisting leukemic cells from the chemosensitive SCXC-7575 PDX did not demonstrate increased basal or max OXPHOS (Fig 4G, Supplemental Fig 4B for all seahorse tracings). Full clinical annotation available in Fig 4H. Altogether, these data suggest that the benefit of a statin is seen when the AML responds to AraC with a mitochondria stress response, but that more robust *in vivo* inhibition of the mevalonate pathway is necessary to improve the clinical response to AraC.

**Figure 4:**
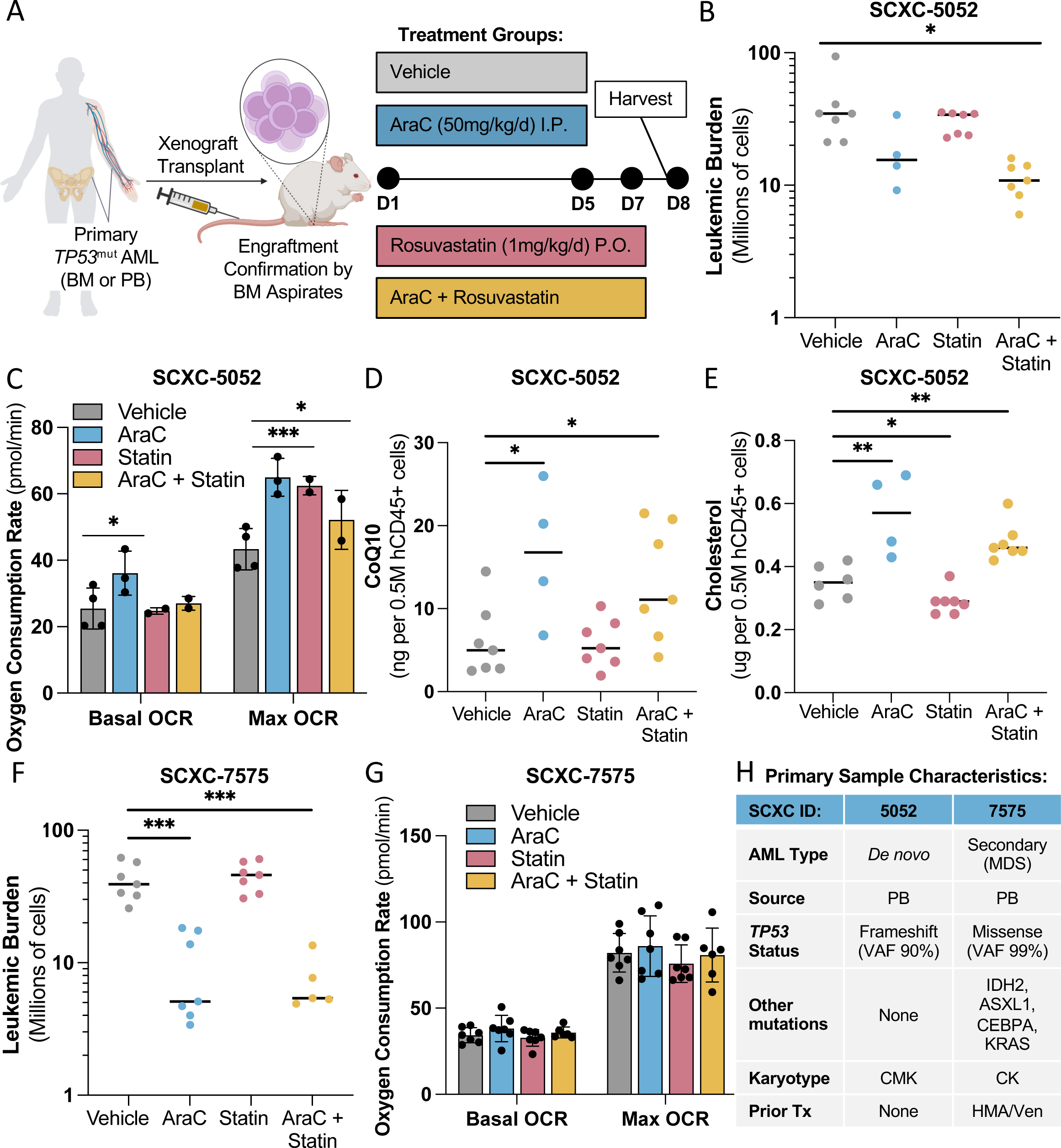
PDX modelling of *TP53*^mut^ AML reveals statin-mediated AraC chemosensitization. (A) Schema (created in BioRender) of *TP53*^mut^ AML PDX model. In brief, primary, human *TP53*^mut^ AML cells were injected into the tail veins of busulfan-pretreated NSG mice. Engraftment was confirmed on bone marrow aspirates followed by initiation of treatment with vehicle, AraC (50mg/kg/day IP for D1-5), rosuvastatin (1mg/kg/day PO for D1-7) or the two drugs in combination. On D8 mice were humanely sacrificed and bilateral femurs/tibias and spleen were harvested (n=7 mice per condition, early death of 3 mice in AraC condition). (B) Leukemic burden in millions of hCD45+ hCD33+ cells (chemoresistant sample SCXC-5052) in the bone marrow and spleen combined for each mouse and quantified with counting beads by flow cytometry. (C) Basal and maximum coupler-induced oxygen consumption in pmol/minute assessed by Seahorse technology in magnetic bead-purified hCD45+ leukemic cells from the bone marrow with each circle representing the average of 1 mouse (n=4, 3, 2, 2 mice for Control, AraC, Statin and AraC + Statin, respectively; chemoresistant sample SCXC-5052). (D) CoQ10 (ng) and (E) cholesterol (ug) per 0.5 million magnetic bead-purified hCD45+ leukemic cells from the bone marrow of mice with each circle representing the average of 1 mouse, as assessed by LC-HRMS (5 replicates per mouse; n=7, 4, 7, 7 mice for Control, AraC, Statin and AraC + Statin, respectively; chemoresistant sample SCXC-5052). (F) Leukemic burden and (G) oxygen consumption rate as described in (B) and (C) but for chemosensitive sample SCXC-7575. (H) Summary of primary sample characteristics.

### *In vitro* statin pretreatment fully abrogates OXPHOS and chemoresistance in primary *TP53*^mut^ AML

To assess the efficacy and mechanism of the drug combination on primary samples without the complexities of *in vivo* statin administration(40–42), we studied 4 primary samples (including SCXC-5052 and SCXC-7575 used for the PDX studies above) *in vitro* and assessed OXPHOS by Seahorse after 24-hour pretreatment with rosuvastatin followed by an additional 24 hours of AraC. Three of the 4 samples demonstrated a significant increase in basal and max OXPHOS in response to AraC [For basal and max OXPHOS, respectively: SCXC-4708 1.4-fold (p < 0.001) and 1.3-fold (p < 0.001); SCXC-6865 1.9-fold (p = .005) and 1.7-fold (p = 0.02); SCXC-5052 1.9-fold (p < 0.001) and 1.7-fold (p < 0.001)], which was blocked by rosuvastatin pretreatment (Fig 5A). Those 3 samples also demonstrated a dose-dependent decrease in colony forming unit ability by the combination of AraC and rosuvastatin (Fig 5B). Consistent with the *in vivo* data, SCXC-7575, did not significantly increase OXPHOS in response to AraC [1.1-fold (p = 0.13) and 1.1-fold (p = 0.2) for basal and max OXPHOS, respectively) (Supplemental Fig 5A) with a smaller benefit of combination treatment on CFUs (Supplemental Fig 5B). Furthermore, two of the three AraC-resistance primary samples responded to AraC with an increase in mitochondria mass as measured by flow cytometry after staining for mitochondria protein, TOMM20 (Supplemental Fig 5C). These studies suggest that primary human *TP53*^mut^ AML cells treated with AraC require the mevalonate pathway for survival via regulation of OXPHOS, which is blocked by rosuvastatin. Similar to our *in vivo* results, there is no additional benefit of rosuvastatin if the primary human AML sample is unable to induce the mitochondria stress response to AraC, validating the OXPHOS-specific mechanism of rosuvastatin. Finally, compared to our *in vivo* studies where rosuvastatin only partially reversed AraC-induced OXPHOS, the statin concentrations achieved *in vitro* were able to fully block AraC-induced OXPHOS in primary samples. We next sought to further define the targetable branches of the pathway to find alternative mechanisms to fully inhibit the mevalonate pathway *in vivo*.

**Figure 5:**
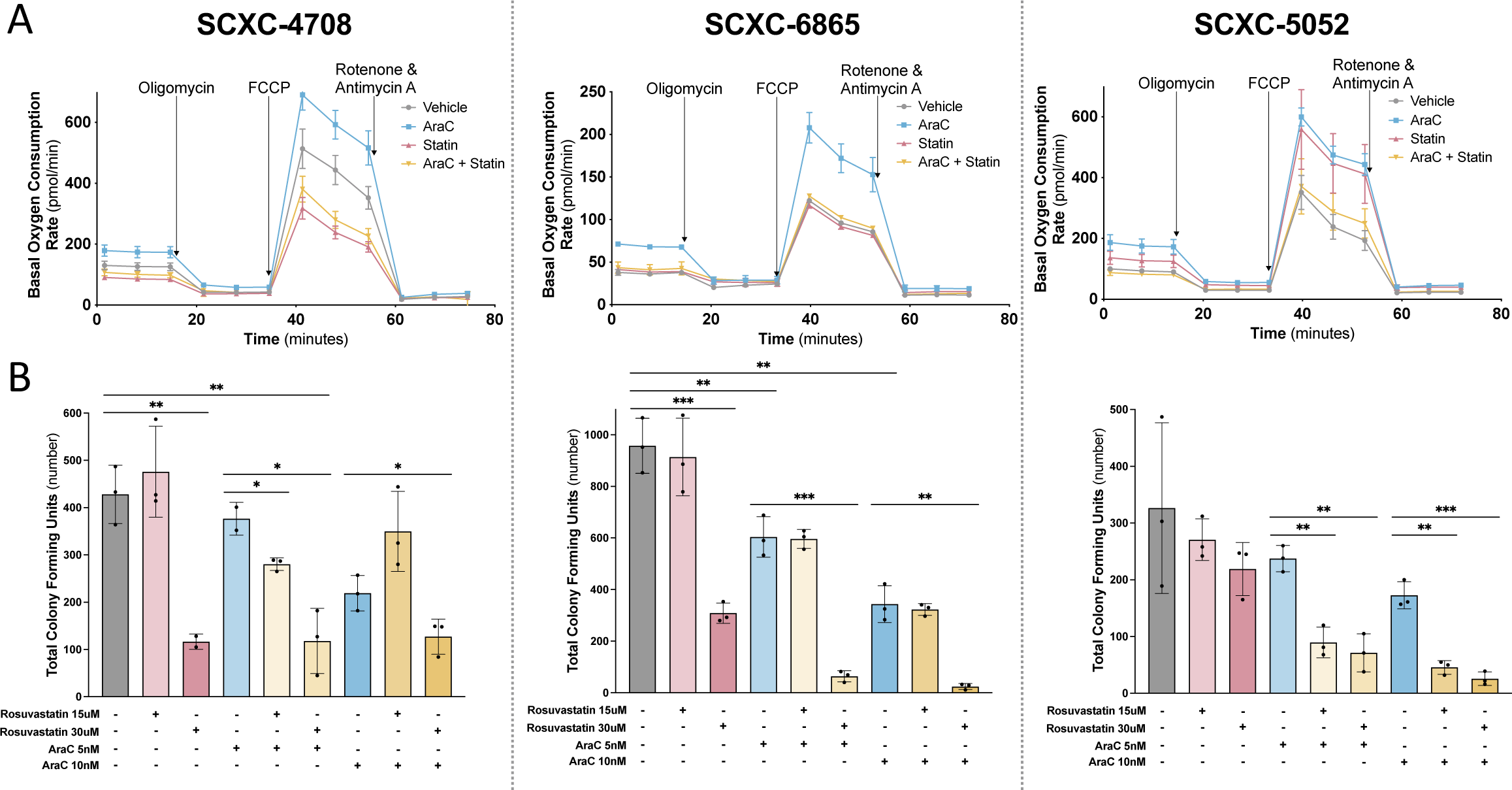
*In vitro* statin pretreatment fully abrogates OXPHOS and chemoresistance in primary *TP53*^mut^ AML. (A) Oxygen consumption rate (pmol/min) assessed by Seahorse technology in previously viably frozen primary *TP53*^mut^ AML samples that had been resuspended in X-Vivo media with 20% BIT serum and 10ng/mL of cytokines FLT3, SCF, IL3 and IL6, pretreated with rosuvastatin (50uM) for 24h followed by AraC (1uM) treatment for an additional 24h, and underwent dead cell depletion *via* the AnnexinV dead cell removal kit prior to plating for Seahorse analysis. Each sample (SCXC-4708, −6865, −5052) was performed once with 5 technical replicates per condition. (B) Total number of colony forming units assessed 14 days after plating previously viably frozen primary *TP53*^mut^ AML samples (SCXC-4708, −6865, −5052) treated on day 0 with rosuvastatin (15uM or 30uM) and/or AraC (5nM or 10nM) with 3 replicates per condition. Statistical analysis by Student’s T Test. p-values: * = <0.05, ** = <0.01, *** = <0.001. n is the number of replicates.

### AraC resistance of *TP53*^mut^ AML requires the mevalonate pathway byproduct, GGPP, for ROS regulation and enhanced OXPHOS

To determine the specific mevalonate pathway byproducts necessary for chemoresistance, we performed rescue experiments in the representative *TP53*^mut^ AML clone, M14-Mut1, treated with a rescue agent and rosuvastatin for 48 hours with AraC added for the last 24 hours (Schema summarizing targets in Fig 6A). Co-treatment of M14-Mut1 with mevalonate (MVA), which is produced by the direct statin target, HMGCR, completely rescued the effect of rosuvastatin on cell viability (Fig 6B) and OXPHOS (Fig 6C, Supplemental Fig 6A-B). Co-treatment of M14-Mut1 with downstream GGPP also rescued the effects of a statin on cell viability (Fig 6D), OXPHOS (Fig 6E, Supplemental Fig 6C-D), TOMM20 as a surrogate for mitochondria mass (Fig 6F), mitochondrial ROS (Fig 6G), and total and reduced glutathione (Fig 6H, Supplemental Fig 6E). GGPP did not rescue the rosuvastatin-induced decrease in coenzyme Q10 in either M14-Mut1 or M14-WT1 (Supplemental Fig 6F), indicating that the CoQ10 isoprenoid tail requires upstream FPP, but not GGPP. Altogether, these data indicate that rosuvastatin’s activity is specific to the mevalonate pathway, independent of cholesterol, and dependent on the GGPP branch of the pathway. The presence of GGPP correlates with improved glutathione synthesis and reduced ROS, specifically in the *TP53*^mut^ AML clones (M14-WT1 experiments in Supplemental Fig7). Finally, we validated that specific geranylgeranyl transferase (GGT) I inhibitor, GGTI298, but not GGTII inhibitor, GGTI2133, or farnesyltransferase (FNT) inhibitor, FTI277, recapitulates rosuvastatin’s impact on cell viability alone or in combination with AraC (Figure 6I, Supplemental Fig 6G-H). Importantly, there is a synergistic decrease in cell viability in M14-Mut1 treated with both rosuvastatin and GGTI298 [F(3,12)=20, p < 0.001)] with a further benefit of the triplet therapy with AraC [F(7,24)=16, p < 0.001)], suggesting that dual mevalonate pathway inhibition may help overcome incomplete inhibition of the mevalonate pathway by rosuvastatin *in vivo* and potentially in patients.

**Figure 6:**
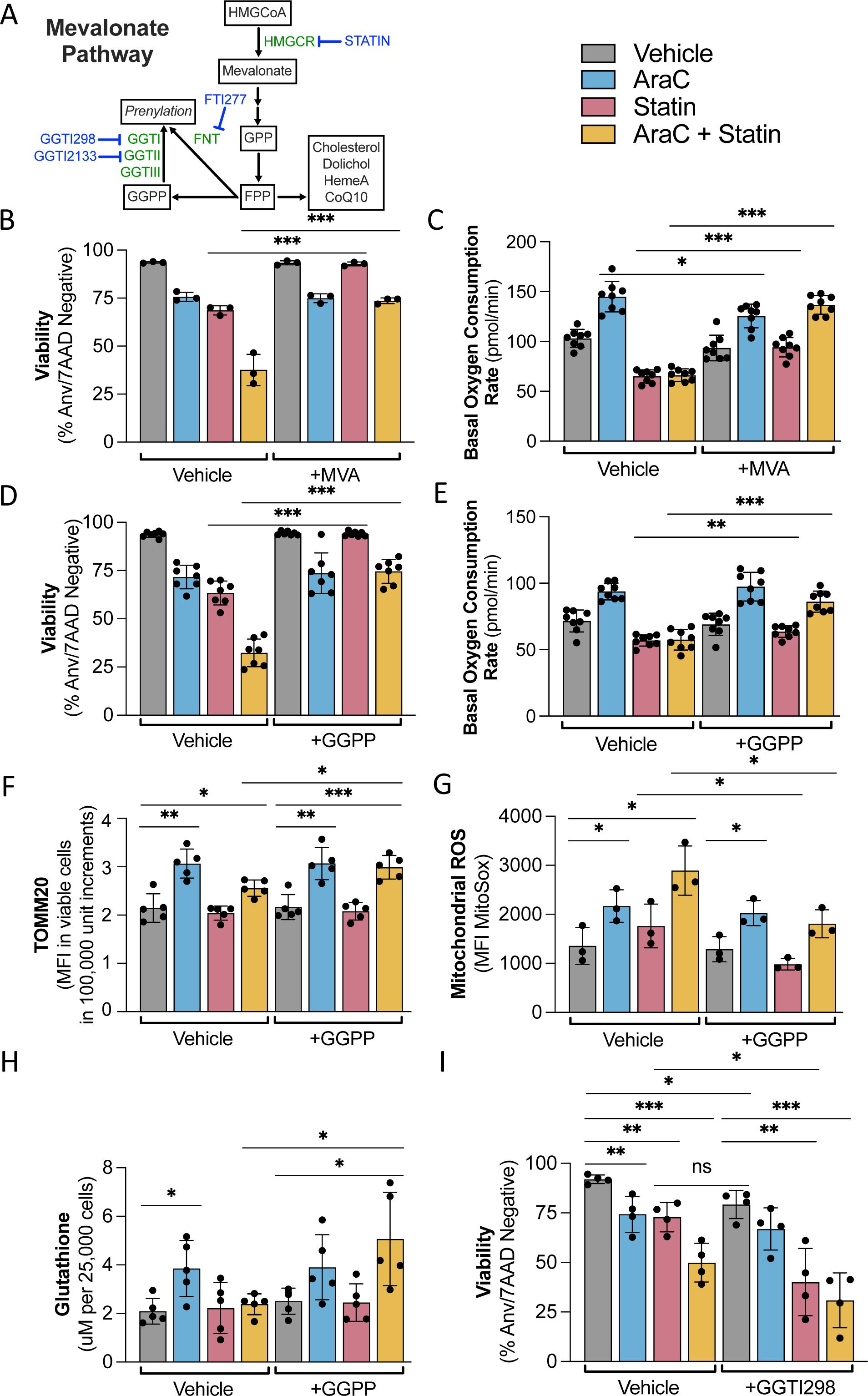
AraC resistance of *TP53*^*m*ut^ AML requires GGPP for ROS regulation and enhanced OXPHOS. (A) Abridged schema of latter half of the mevalonate pathway with essential genes in green and known chemical inhibitors in blue. All experiments in Figures 6B-I were performed in the M14-Mut1 clone pretreated for 24h with rosuvastatin (50uM) and either vehicle, MVA (200uM), GGPP (1uM) or GGTI298 (10uM) followed by an additional 24h with vehicle or AraC (1uM) with the subsequent assessment of (B) cell viability presented as the percentage of annexinV and 7AAD negative cells by flow cytometry with or without MVA (n=3), (C) basal oxygen consumption rate in pmol/minute as assessed by Seahorse technology with or without MVA (n=8), (D) cell viability presented as the percentage of annexinV and 7AAD negative cells by flow cytometry with or without GGPP (n=7), (E) basal oxygen consumption rate in pmol/minute as assessed by Seahorse technology with or without GGPP (n=8), (F) TOMM20 presented as MFI in viable cells (by FVS-R) assessed by flow cytometry with or without GGPP (n=5), (G) mitochondrial ROS presented as the ratio to vehicle of MFI of MitoSox in viable cells (by FVS-R) as measured by flow cytometry with or without GGPP (n=3), (H) total glutathione presented as uM per 25,000 cells with or without GGPP (n=5), and (I) cell viability presented as the percentage of annexinV and 7AAD negative cells by flow cytometry with or without GGTI298 (n=4). Statistical analysis by Student’s T Test. p-values: * = <0.05, ** = <0.01, *** = <0.001. n is the number of replicates.

## Discussion

*TP53* mutations are notorious for conferring chemoresistance in AML, rendering standard treatments largely ineffective and leading to dismal patient outcomes(1–3). Our study delves into the mechanistic insights and therapeutic implications of targeting the mevalonate pathway in *TP53*^mut^ AML. We demonstrate for the first time that *TP53*^mut^ AML requires a mevalonate pathway-dependent mitochondria stress response to survive chemotherapy. Furthermore, we provide compelling evidence that the GGPP branch of the pathway is crucial for mitochondria-dependent chemoresistance potentially through its regulation of mitochondria biogenesis and a novel contribution to glutathione synthesis for management of chemotherapy-induced ROS.

Our group and others have previously shown mutation-agnostic AML dependence on mitochondria metabolism(6,43,44), particularly in response to chemotherapy(5,7–9). A robust study evaluating the impact of *TP53* missense mutations in solid tumor malignancies, such as non-small cell lung cancer, hepatocellular carcinoma and breast adenocarcinoma, found that stable endogenous *TP53* mutations led to *decreased* OXPHOS whereas inducible and transient models had *increased* OXPHOS(45). Our data supports these findings in which stable *TP53*^mut^ AML single cell clones at baseline have *decreased* OXPHOS (Fig 3A-B, Supplemental Fig 3A-B). However, when the cells are stressed with AraC, for example, we see a significant *increase* in OXPHOS that may be similar to the inducible/transient *TP53*^mut^ solid tumor models. We recognize our model is different from a model of p53 loss in the AML cell line, MOLM-13, in which clones developed with two different guide RNAs targeting *TP53* had increased OXPHOS at baseline(10). It is crucial to evaluate OXPHOS specifically in the context of stress in *TP53*^mut^ vs *TP53*^WT^ tumor models to understand the impact of p53 alterations on mitochondria metabolism and subsequent therapeutic responses. Importantly, this finding also indicates *TP53*^mut^ AML shares common mechanisms of chemoresistance with refractory *TP53*^WT^ AMLs described in the literature(5,7–9). We propose that *TP53*^mut^ AML is the archetype of refractory AMLs and that studying chemoresistance in this context will translate to other therapy-resistant leukemias.

Our study also furthers our understanding of the mechanism of action of AraC. Using novel isogenic *TP53*^mut^ AML cell lines, we observe that the ability to induce the mitochondria stress response in *TP53*^mut^ AML correlates with glutathione antioxidant capacity and the ability to regulate AraC-induced ROS. Conversely, the *TP53*^WT^ AML cells exhibit a rapid and significant increase in ROS in response to AraC, despite no significant increase in OXPHOS. This paradoxical finding demonstrates that increased OXPHOS can be associated with decreased ROS, which we propose is due to more efficient ROS management particularly through upregulation of glutathione synthesis for its role as an antioxidant. Our data also shows that AraC-induced ROS may be one of the critical events that leads to its toxicity. Traditionally, the increase in ROS is thought to be due to genomic DNA damage(46). However, AraC is incorporated into DNA in the S-phase of the cell cycle. Given our cell lines have a doubling time of 24 hours and patient samples are not necessarily proliferating *in vitro*, genomic DNA damage as the primary mechanism of ROS induction appears unlikely. Alternatively, we suggest mitochondria DNA damage(47) or nucleotide imbalance(48) may lead to an increase in ROS and a subsequent need for a mitochondria stress response, including increased mitochondria biogenesis, to survive.

Direct mitochondria targeting to reverse chemoresistance remains a challenge in the field(13). Inhibiting the mevalonate pathway may be an indirect but effective way to overcome toxicity from direct mitochondria inhibition. In fact, particularly in our primary sample models *in vitro* and *in vivo*, only AraC-resistant *TP53*^mut^ AML samples that induce the mitochondria stress response are sensitive to concurrent mevalonate pathway inhibition during AML therapy. This suggests a therapeutic window in which mevalonate pathway-addicted cells will be most sensitive to pathway inhibition. Our retrospective analysis shows that a broad range of statins at different doses appear to be well tolerated and safe in combination with AML therapy in *TP53*^mut^ AML patients (Fig 1D). Furthermore, there appears to be an early survival benefit of statins in the first 6 months following *TP53*^mut^ AML diagnosis. However, statin efficacy in mouse and human models remains limited by (1) statin pharmacokinetics and pharmacodynamics that make it challenging to achieve preclinical doses in humans(14,40), which is further exacerbated in rodent models(42), (2) rescue of mevalonate pathway inhibition by dietary presence of mevalonate byproducts(41), and (3) highly conserved feedback loops that induce mevalonate pathway upregulation following inhibition(38).

To optimize mevalonate pathway inhibition for anti-cancer benefit, it is essential to understand the crucial role of the mevalonate pathway in chemoresistance and identify mechanisms to improve targeting in patients. Through extensive rescue studies and chemical validation, we determined that GGPP is the essential byproduct downstream of mevalonate pathway inhibition that leads to toxicity of rosuvastatin as a single agent and in combination with AraC. Other groups have validated the importance of this byproduct in multiple liquid and solid tumor models(24,25,27,29,49,50). However, the specific function of GGPP has remained elusive likely due to GGPP’s role in regulating many GTPases and that the essential function of GGPP at a given time may be cell-type and cell-context specific. In our studies, GGPP may have multiple roles, such as (1) regulation of GTPases that are crucial for mitochondria biogenesis and (2) contributing to intracellular glutathione levels. Notably, GGPP-dependency for maintaining glutathione has only been previously described in adipose tissue(51) and the specific mechanism by which GGPP regulates glutathione remains unknown. Glutathione synthesis appears to be particularly important for chemoresistance as only isogenic *TP53*^mut^ AML cell line clones can significantly increase total and reduced glutathione in response to AraC, which is blocked by rosuvastatin in a GGPP-dependent manner. Future studies will define the specific mechanisms by which GGPP regulates mitochondria biogenesis and the intracellular glutathione pool, and determine the therapeutic benefit of direct targeting of GGPP in *TP53*^mut^ AML.

## Methods

### Cell culture and reagents

Human AML cell lines (Supplemental Table 1F) were maintained in minimum essential medium-alpha (Gibco) supplemented with 10% fetal bovine serum (Hyclone) at 37°C and 5% CO2. The cultured cells were split every 2 to 3 days and maintained in an exponential growth phase. Authenticated MOLM14 cells were obtained from DSMZ and the liquid nitrogen stock was renewed every 2 years. Cell lines were routinely tested for mycoplasma contamination. Cytarabine (Hospira) was obtained from The Hospital of The University of Pennsylvania and diluted in H2O for use *in vitro*. Rosuvastatin (Sigma), pitavastatin (MedChemExpress), GGTI298 (Sigma), GGTI2133 (Sigma), and FTI277 (Sigma) were dissolved in DMSO for use *in vitro*. Reduced glutathione (Sigma) was directly dissolved in media and buffered with NaOH (Sigma). Mevalonolactone (Sigma) is dissolved in ethanol. Geranylgeranyl pyrophosphate (Sigma) is suspended in methanol. For all experiments, drug is diluted so that final vehicle concentration is 0.1% for all experiments.

### Primary AML specimens

Mononuclear cells (MNCs) from patients with AML were obtained from the Stem Cell and Xenograft Core (SCXC, IRB Protocol Number: 703185) at the University of Pennsylvania. All samples were acquired through the SCXC following informed consent on the institutional review board-approved protocol. Samples were collected in accordance with federal and institutional guidelines and provided to us in a pathologically annotated and de-identified fashion.

### Flow sorting

Patient samples were thawed and washed with PBS supplemented with 5% FBS (Corning) supplemented with DNase I (Sigma) at 50ug/mL. Cells were incubated with a fixable viability stain 510 (BD Biosciences) for 15 min at room temperature, washed, and then stained with antibodies against human CD45 (BD Horizon BV786), CD33 (BD Horizon BV421), CD3 (BD Horizon APCR700), and CD19 (BD Horizon BB515) for 20 min at room temperature. Cells were then washed and resuspended in PBS supplemented with 2% FBS and flow sorted to isolate viable (FVS510^−^) CD3^−^/CD19^−^/CD45+ / CD33+ cells using a BD FACSAria III cell sorter.

### RNA-Sequencing Analysis

Following flow sorting, primary samples were washed in PBS and then extraction was performed using the Qiagen AllPrep DNA/RNA kit. The purity and concentration of the extracted DNA and RNA were assessed using a Nanodrop before downstream applications. RNA was sent to Tempus for library preparation and sequencing on an Illumina HiSeq. RNA libraries were prepared from PolyA selected mRNA species. Multiplexed sequencing was performed with paired-ends to a read depth of 20-30 million per sample. FASTQ files were aligned to hg19 using STAR Align version 2.7.8a(52). Genes were counted using HTSeq-Count version 2.0.3(53). Gene expression data in the form of counts per million was log2 transformed with a pseudocount of 0.5 added to all values and z-scored. Single sample gene set enrichment analysis was performed using the GSVA package in R. Gene signatures were tested from the MsigDB C2 or Hallmark collection. Statistical differences between *TP53*^mut^ and *TP53*^WT^ patients were assessed by Student’s T-test. RNA sequencing analysis was also performed on *de novo* AML patients from the BeatAML cohort(54) obtained from the public database(54).

### Retrospective clinical data

364 total patients diagnosed with *TP53*^mut^ AML were evaluated in this study, including 215 treated at The University of Pennsylvania and 149 from Roswell Park Cancer Center. Both institutions consent patients independently for data collection for an electronic health record-derived database. Using the databases, we identified patients diagnosed with AML between 2013 and 2023 with a *TP53* mutation at the time of AML diagnosis. We performed chart reviews to determine treatment with a statin before and/or during initial treatment for AML as well as baseline patient demographics and clinical outcomes. Kaplan-Meier methodology was performed for median overall survival.

### Generation of isogenic *TP53*^mut^ AML cell lines

sgRNA sequences for negative control (Rosa) and TP53 targeting (Gia 5 and Grum A) are as follows: GAAGATGGGCGGGAGTCTTC, CATGTGTAACAGTTCCTGCA and GGGCAGCTACGGTTTCCGTC, respectively(35,36). These were ligated into LentiCRISPRv2-mCherry (Addgene #99154), digested with BsmBI (NEB #R0580). Lentivirus was packaged in 293T cells cotransfected with pHEF-VSVG (Addgene#22501) and pPax2 (Addgene#12259) in PEI transfection reagent. Lentivirus supernatant was harvested at 2- and 3-days post transfection and was concentrated with PegIT Precipitation (SBI #LV810A-1). MOLM14 cells were then incubated with virus overnight and transduction was verified by flow cytometry for mCherry. Mutations were validated by DNA sequencing and loss of TP53 function confirmed by Western blotting after AraC, as described above. Single cell clones were selected by serial dilution and expanded. Loss of function of p53 was confirmed again in all single cell clones.

### DNA sequencing

Cell lines were washed twice with PBS and flash frozen as pellets. Subsequently, genomic DNA was isolated using a the Qiagen DNeasy Blood and Tissue Kit. DNA from primary samples was extracted with the Qiagen AllPrep DNA/RNA kit as described above. Libraries of germline DNA were prepared using KAPA Hyper Prep Kit (Roche Diagnostics, Branchburg NJ). Libraries were subjected to targeted next generation sequencing (NGS) of the *TP53* gene on an Illumina Nova-Seq. FASTQ files from sequencing were aligned to human genome version 38 (hg38) using BWA-mem version 0.7.17(55). Germline variants were called from BAM files using Genome Analysis Toolkits (GATK) HaplotypeCaller version 3.7. The presence of *TP53* variants was validated manually using the Integrative Genomics Viewer version 2.5.2.

### Apoptosis assay

Unfixed cells were resuspended in Annexin Binding Buffer (10 mM HEPES, 140 mM NaCl, 2.5 mM CaCl2 pH 7.4) and stained with Annexin V-APC (Invitrogen) and 7AAD (BD Biosciences). Stained cells were analyzed using a BD Accuri C6 flow cytometer. Data was processed with FlowJo version 10 (BD Biosciences).

### Immunoblot

Cells were lysed with the CellLytic MT Cell Lysis Reagent (Sigma) with protease and phosphatase inhibitors (Halt Protease & Phosphatase Inhibitor Cocktail; Thermo Fisher). Lysates were analyzed by SDS-PAGE. Immunoblot detection was performed using Amersham ECL Detection Reagents (Cytiva). Antibodies used are as follows: p53 (EMD Millipore #OP43), p21 (BD Pharmingen #556430), HMGCR (Invitrogen #MA5-35242), SREBP2 (Invitrogen #PA1-338), TBP (Cell Signaling Technologies [CST], #44059). Additional reagents used include a SREBP2 positive control lysate (Origene #LC417881) and secondary antibodies (CST #7074, CST #7076). Signal intensity was quantified using ImageJ.

### XTT Cell viability assay

Cells were pre-treated with rosuvastatin or DMSO for 24 hours. Cells were then plated in 96-well plates at 100,000 per well in triplicate with AraC plus fresh DMSO or rosuvastatin. Cell viability was measured after 24 hours of AraC treatment using the XTT Cell Viability Kit from Cell Signaling Technology (#9095) and a Tecan Infinite 200 Pro plate reader. Drug-drug interaction was evaluated using the SynergyFinder web-based tool, with a score greater than 10 being likely synergistic, −10 to 10 likely additive, and less than −10 likely antagonistic(37).

### Quantitative reverse transcription polymerase chain reaction (qRT-PCR)

AML cell lines were harvested, washed with PBS, and flash frozen in liquid nitrogen as cell pellets. RNA extraction was performed using the QIAGEN RNeasy Mini Kit following the manufacturer’s protocol with the following adjustments. Cells were lysed using Buffer RLT supplemented with DNase I to remove genomic DNA and homogenized using QIAshredders. The quantity and quality of isolated RNA was assessed with the Nanodrop One. Total RNA extracted was then reverse transcribed into complementary DNA (cDNA) utilizing the BIO-RAD iScript RT Supermix according to the manufacturer’s instructions. qRT-PCR was performed using TaqMan Fast Advanced Master Mix and predesigned TaqMan probes on a 384-well plate using the ViiA 7 Rea-time PCR system (Thermo Fisher). The predesigned TaqMan probes used are as follows: ABCA1 (Hs00194045), SREBF1 (Hs02561944), SREBF2 (Hs01081784), HMGCR (Hs00168352), IDI1 (Hs00743568), TRIT1 (Hs01091215), FDPS (Hs01578769), GGPS1 (Hs01546492), FNTA (Hs00357739), PGGT1B (Hs00270701), RABGGTA (Hs01554344), RABGGTB (HS00190183), PDSS1 (Hs00372008), PDSS2 (Hs01047689), FDFT1 (Hs00926054), SQLE (Hs01123768), LSS (Hs01552331), GAPDH (Hs02786624), CDKN1A (Hs00355782), MYC (Hs00153408).

### Liquid chromatography-high resolution mass spectrometry (LC-HRMS)

Cells are resuspended in fresh, pre-warmed media at 400,000 cells per mL and treated as described, with 5 replicates per condition. At the end of treatment, cells were counted and 2 million cells per condition were washed twice in cold PBS and flash frozen as cell pellets. The cell pellets were then spiked with a panel of stable isotope labeled lipids as internal standards for normalization of extraction and analysis. The lipid extraction and LC-HRMS analysis was done as previously described(56).

### Seahorse Assay

All XF assays were performed using the Agilent Seahorse XFe96 Extracellular Flux Analyzer. The day before the assay, the sensor cartridge was placed into the calibration buffer medium supplied by Seahorse Biosciences to hydrate overnight. Seahorse XFe96 microplates wells were coated with 25 μl of Cell-Tak (Corning; Cat#354240) solution at a concentration of 22.4 μg/ml at room temperature for 20 minutes, washed twice with distilled water, and kept at 4°C overnight. On the day of the experiment, AML cells were plated at a density of 80,000 cells per well for cell lines and 100,000 cells per well for primary samples in XF base minimal DMEM media containing 11 mM glucose, 1 mM pyruvate and 2 mM glutamine. Then 180 μl of XF base minimal DMEM medium was added to each well and the microplate was centrifuged at 100 g for 1 min with no break. After no more than one hour of incubation at 37°C in CO2 free-atmosphere, basal oxygen consumption rate (OCR, as a mitochondria respiration indicator) and extracellular acidification rate (ECAR, as a glycolysis indicator) were performed using the Seahorse XF Cell Mito Stress Test Kit (#103015-100).

### Mitochondria DNA quantification

Genomic DNA was extracted using the Qiagen DNeasy Blood and Tissue Kit. The purity and concentration of the extracted DNA was assessed using a Nanodrop before downstream applications. PCR was performed using the PowerTrack SYBR Green Master Mix and the primers listed below to determine the ratio of mitochondria to nuclear DNA using a Thermo Fisher Scientific ViiA 7 Real-Time PCR System. Mitochondria DNA (*ND2)*: forward primer sequence – CCTATCACCCTTGCCATCAT; reverse primer sequence – GAGGCTGTTGCTTGTGTGAC. Nuclear DNA (*Pecam1*): forward primer sequence – ATGGAAAGCCTGCCATCATG; reverse primer sequence – TCCTTGTTGTTCAGCATCAC.

### Measurement of ROS content and mitochondria mass

Mitochondrial ROS content and mass in viable cells were measured by flow cytometry using MitoSOX Red Mitochondria Superoxide Indicator (Thermo Fisher) and Tom20 Antibody (5-10) Alexa Fluor488 (Santa Cruz).

### Glutathione Assay

Total glutathione and the ratio of reduced to oxidized glutathione (GSH/GSSG) were determined using the GSH-Glo Glutathione Assay from Promega, which is a luminescence-based assay. Assay was performed and analyzed by their protocol using 25,000 cells per well in triplicate and analyzed with a Tecan Infinite 200 Pro plate reader.

### Colony-forming assays

Primary human AML cells were plated in cytokine-enriched methylcellulose (R&D Systems HSC005) in triplicate. Plates were maintained at 37^º^C and 5% CO2. Drug treatments were added only at the time of plating. Human mononuclear cells were seeded at 10,000-150,000 cells per plate for AML. Colonies were scored at 14 days. Only data from samples with growth of >30 colonies per plate in the DMSO control condition were included.

### Patient-derived xenograft model

Animals were used in accordance with a protocol reviewed and approved by the Institutional Animal Care and Use Committee at UPENN. NOD/LtSz-SCID/IL-2Rγchain^null^ (NSG) mice were produced at The Stem Cell and Xenograft Core using breeders obtained from The Jackson Laboratory. Mice were housed and human primary AML cells were transplanted as reported previously(5). Briefly, mice were housed in sterile conditions using HEPA-filtered microisolators and fed with irradiated food and sterile water. Mice (6–9 weeks old) were sublethally treated with busulfan (30 mg/kg/day) 24 hours before injection of leukemic cells. Leukemia samples were thawed at room temperature, washed twice in PBS, and suspended in PBS at a final concentration of 10-20 million cells per mL. 100 μL (1-2 million cells) was injected into the tail vein of each mouse. Daily monitoring of mice for symptoms of disease (ruffled coat, hunched back, weakness, and reduced mobility) determined the time of killing for injected animals with signs of distress. If no signs of distress were seen, mice were initially analyzed for engraftment 8 weeks after injection except where otherwise noted.

### *In Vivo* drug administration

Four to 12 weeks after AML cell transplantation and when mice were engrafted (confirmed by flow cytometry on bone marrow aspirates), mice were treated by daily intraperitoneal injection of AraC (Hospital of The University of Pennsylvania) 50 mg/kg for 5 days or Vehicle (PBS) or daily oral administration of rosuvastatin (Sigma) 1mg/kg for 7 days or Vehicle (4% DMSO, 30% PEG-300, double distilled H2O). Mice were monitored for toxicity and provided nutritional supplements as needed.

### Assessment of leukemic engraftment

Mice were harvested on days 8 and 9 following initiation of drug treatment. NSG mice were humanely killed in accordance with ethics protocols. Bone marrow (mixed from tibias and femurs) and spleen were dissected in a sterile environment and crushed in 15mL PBS with 2% FBS and filtered through 40uM filters. Samples were centrifuged and resuspended in 2mL PBS with 2% FBS. 100uL per sample was removed for assessment of leukemic engraftment by flow cytometry. 50uL antibody mix was added and cells were incubated for 20 minutes in the dark at room temperature. Antibody mix was composed of human CD45-PE (BD Pharmingen), mouse CD45-PE-Cy7 (BD Pharmingen), CD33-BB515 (BD Horizon), CD3-BV421 (BD Horizon) and CD19-BB700 (BD Horizon). 750uL of 1X BD FACS Lysing Solution was added to each tube and incubated for 15 minutes at room temperature. Cells were washed and resuspended in 300uL of 2% paraformaldehyde. 20uL of CountBright Absolute Counting Beads (Invitrogen) were added and immediately vortexed. Analyses were performed on a Life Science Research II (LSR II) flow cytometer with DIVA software (BD Biosciences). The number of AML cells/μL bone marrow or spleen were determined by using the CountBright beads protocol (Invitrogen).

### Magnetic bead separation and downstream processing of leukemic cells in the PDX model

Following 100uL removal for flow cytometry, 1.9mL of remaining BM sample was centrifuged and then lysed with ammonium chloride for 5 minutes. Cells were washed, centrifuged and then processed for magnetic bead separation using human CD45 microbeads (Miltenyi) *via* their protocol using the Miltenyi autoMACS. Following cell harvest, cells were resuspended in PBS with 2% FBS and divided for flow cytometry sort purity determination, LC-HRMS or Seahorse. For LC-HRMS, 3-5 replicates 0f 0.5 million cells per mouse were washed and then flash frozen as cell pellets and stored in a −80C freezer. For Seahorse, cells were resuspended at 1 million cells per mL in IMDM supplemented with 20% FBS. After 1 hour incubation at 37°C and 5% CO2 cells were processed for Seahorse analysis per above.

### Statistics

All statistics not otherwise described in the methods or figure legends were performed using two-tailed Student’s T test in Prism 10 (GraphPad). The number of independent replicates used for statistical calculations are noted in the legends. Asterisks denote statistical significance versus control condition unless otherwise indicated (*p < 0.05, **p < 0.01, ***p < 0.001).

### Data Availability Statement

The sequencing data will be made available on the GEO database at final publication. All code is available at https://github.com/bowmanr/Skuli_TP53_mevalonate.

## Acknowledgements

We would like to acknowledge the UPenn Stem Cell and Xenograft Core for their excellent support for all *in vivo* experiments and tissue banking. Research support for this study was generously provided by The UPenn Hematology Clinical Research Training Program through the National Institutes of Health (T32HL0439; 2020-2022, SJS); American Society of Hematology Research Training Award for Fellows (2022-2022, SJS); American Society of Clinical Oncology Young Investigator Award (2023-2024, SJS), UPenn Clinical and Translational Research Award KL2 Program through the NIH (KL2TR001879; 2023-2025, SJS); Department of Defense Impact Award (W81XWH-20-1-0867, DAF); National Cancer Institute of the NIH (P30ES013508, CM; R35CA220483, MCS; K08CA230190, PJS; P30CA16056, PJS, LNM-G; U54CA283759, MC); a Veterans Merit Award (I01BX000918, MC); The Laboratoire d’Excellence Toulouse Cancer (ANR11-LABEX, TOUCAN and TOUCAN2.0; J-ES); INCA (HRHG-MP22-036, IRONAML53; J-ES); the Ligue Nationale de Lutte Contre le Cancer (J-ES); the association GAEL (J-ES). The content is solely the responsibility of the authors and does not necessarily represent the official views of the NIH, NCI or foundations. J-ES is a member of OPALE Carnot Institute and The Organization for Partnerships in Leukemia.

## Notes

**Disclosure of Potential Conflicts of Interest:** CL is a consultant for Bristol Myers Squibb 54 (BMS), AbbVie and Astellas; CL is on advisory boards for Servier, Daiichi, AbbVie, Rigel and Genentech; CL receives research funding from JAZZ and BMS. DAF is a consultant for AbbVie. Otherwise none.

https://github.com/bowmanr/Skuli_TP53_mevalonate

### Competing Interest Statement

CL is a consultant for Bristol Myers Squibb (BMS), AbbVie and Astellas; CL is on advisory boards for Servier, Daiichi, AbbVie, Rigel and Genentech; CL receives research funding from JAZZ and BMS. DAF is a consultant for AbbVie.

### Summary of Updates

This version of the manuscript has been revised to update the disclosures.

