## Supplemental Figures for "Chemoresistance of *TP53* mutant AML requires the mevalonate byproduct, GGPP, for regulation of ROS and induction of a mitochondria stress response"

**Supplemental Figure Legends**

**Supplemental Figure 1:** Kaplan-Meier survival estimates for the primary RNA sequencing (A) UPenn and (B) BeatAML cohorts. RNA sequencing from a cohort of the BeatAML dataset comparing *de novo*, untreated primary $TP53^{mut}$ (n=17) *versus* $TP53^{WT}$ (n=85) AML was analyzed for (C) GSEA scores from the Hallmark and other denoted gene sets, and (D) single sample GSEA with calculation of normalized signature scores for "TP53 Targets," "Hallmark Cholesterol," and "Maxwell Cholesterol." Statistical analysis by Student's T Test.

**Supplemental Figure 2:** (A) Protein expression of p53 and p21 normalized to TBP in representative isogenic $TP53^{mut}$ and $TP53^{WT}$ AML clones treated for 0, 6 or 24 hours with AraC (1uM) performed by western blot. (B) Dose response matrix of M14-WT1 (left) and M14-Mut1 (right) pretreated with 24h of rosuvastatin (0, 25, 50 or 100uM) followed by an additional 24h of AraC (0, 0.5, 1, 2, 4, 8, 16 uM) and assessed by the XTT assay and the Bliss mean synergy score calculator (n=3). (C) Cell viability of isogenic $TP53^{mut}$ and $TP53^{WT}$ AML clones treated for 24 hours with pitavastatin (0, 0.1, 0.3, 1, 3, 10 uM) followed by an additional 24 hours of vehicle or AraC (1uM) and assessed by flow cytometry following staining with AnnexinV and 7AAD. (D) Gene expression of mevalonate pathway genes in isogenic $TP53^{mut}$ and $TP53^{WT}$ AML clones at 0, 6, 12, 18, and 24-hours with AraC (1uM) with all AraC conditions compared to the 0 hour timepoint for each clone after normalization to GAPDH and as measured by qRT-PCR (n=3). (E) ImageJ quantification of HMGCR protein expression normalized to TBP presented as relative units (R.U.) for the western blot in Fig 2E. (F) Protein expression of SREBP2 normalized to TBP in representative isogenic $TP53^{mut}$ and $TP53^{WT}$ AML clones treated for 18h with vehicle or AraC (1uM) performed by western blot with SREBP2+ overexpressed 293T cells as a positive control. Statistical analysis by Student's T Test. p-values: * = <0.05, ** = <0.01, *** = <0.001. n is the number of replicates.

**Supplemental Figure 3:** (A) Oxygen consumption in pmol per minute in M14-Mut2, M14-Mut3, and M14-Mut4 treated with vehicle or AraC (1uM) for 24 hours as measured by Seahorse technology. (B) Summary of basal (left) and maximum uncoupler-induced (right) oxygen consumption in pmol per minute measured by Seahorse technology in the isogenic $TP53^{mut}$ and $TP53^{WT}$ AML clones treated with vehicle or AraC (1uM) for 24 hours. (C) Seahorse tracings for M14-WT1 (left) and M14-Mut1 (right) for Fig 3F. For D-E, representative clones were treated for a total of 48 hours with rosuvastatin (50uM) with AraC (1uM) added for the last 24 hours, and the following experiments were performed: (D) Maximum uncoupler-induced oxygen consumption rate in pmol per minute assessed by Seahorse technology (n=7), (D) ratio of reduced to oxidized glutathione (GSH/GSSG) assessed in 25,000 cells per replicate (n=4). Statistical analysis by Student's T Test. p-values: * = <0.05, ** = <0.01, *** = <0.001. n is the number of replicates.

**Supplemental Figure 4:** Oxygen consumption rate tracings in pmol/minute for (A) Fig 4C and (B) Fig 4G. Top panel includes all seahorse tracings for each PDX experiment, with each line representing 1 mouse. Subsequent panels focus specifically on vehicle *versus* AraC, Statin, and AraC with Rosuvastatin, respectively.

**Supplemental Figure 5:** (A) Oxygen consumption rate (pmol/min) assessed by Seahorse technology in a previously viably frozen primary $TP53^{mut}$ AML sample (SCXC-7575) that had been resuspended in X-Vivo media with 20% BIT serum and 10ng/mL of cytokines FLT3, SCF, IL3 and IL6, pretreated with rosuvastatin (50uM) for 24 hours followed by AraC (1uM) treatment for an additional 24 hours, and underwent dead cell depletion *via* the AnnexinV dead cell removal kit prior to plating for Seahorse analysis. 5 technical replicates per condition. (B) Total number of colony forming units assessed 14 days after plating previously viably frozen primary $TP53^{mut}$ AML samples (SCXC-7575) treated on day 0 with rosuvastatin (15uM or 30uM) and/or AraC (5nM or 10nM) with 3 replicates per condition. (C) TOMM20 presented as MFI in viable cells (by FVS-R) assessed by flow cytometry in previously viable samples (SCXC-4708, -6865, -5052, -7575) after conditioning and treating as per (A) with 1 replicate per condition. Statistical analysis by Student's T Test. p-values: * = <0.05, ** = <0.01, *** = <0.001. n is the number of replicates.

**Supplemental Figure 6:** All experiments in Supplemental Figures 6A-E were performed in the M14-Mut1 clone pretreated for 24 hours with rosuvastatin (50uM) and either vehicle, MVA (200uM) or GGPP (1uM) followed by an additional 24 hours with vehicle or AraC (1uM) and the subsequent assessment of (A) maximum uncoupler-induced oxygen consumption rate in pmol/minute as assessed by Seahorse technology with or without MVA (n=8), (B) representative seahorse tracings presented as oxygen consumption rate (pmol/min) for Fig 6C and Supplemental Fig 6A, (C) maximum uncoupler-induced oxygen consumption rate in pmol/minute as assessed by Seahorse technology with or without GGPP (n=8), (D) representative seahorse tracings presented as oxygen consumption rate (pmol/min) for Fig 6E and Supplemental Fig 6C, and (E) ratio of reduced to oxidized glutathione (GSH/GSSG) assessed in 25,000 cells treated with or without GGPP (n=5). (D) CoQ10 (ng) per 2 million cells were assessed by LC-HRMS in either M14-WT1 or M14-Mut1 cells pretreated for 24 hours with vehicle or rosuvastatin (50uM) and either vehicle or GGPP (1uM) followed by an additional 24 hours with vehicle or AraC (1uM) (n=5). (G) Cell viability as percentage of annexinV and 7AAD negative cells by flow cytometry of M14-Mut1 cells pretreated for 24 hours with vehicle or rosuvastatin (50uM) and either vehicle or GGTI2133 (10uM) followed by an additional 24 hours with vehicle or AraC (1uM) (n=3). (F) Cell viability as percentage of annexinV and 7AAD negative cells by flow cytometry of M14-Mut1 cells pretreated for 24 hours with vehicle or rosuvastatin (50uM) and either vehicle or FTI277 (10uM) followed by an additional 24 hours with vehicle or AraC (1uM) (n=3).

**Supplemental Figure 7:** All experiments were performed in the M14-WT1 clone pretreated for 24 hours with vehicle or rosuvastatin (50uM) and either vehicle, MVA (200uM), GGPP (1uM), GGTI298 (10uM), GGTI2133 (10uM) or FTI277 (10uM) (as indicated in figure) followed by an additional 24 hours with vehicle or AraC (1uM) with the subsequent assessment of (A) cell viability as the percentage of annexinV and 7AAD negative cells by flow cytometry with or without MVA (n=3), (B) basal oxygen consumption rate in pmol/minute as assessed by Seahorse technology with or without MVA (n=8), (C) maximum uncoupler-induced oxygen consumption rate in pmol/minute as assessed by Seahorse technology with or without MVA (n=8), (D) representative Seahorse tracings presented as oxygen consumption rate (pmol/min) for Supplemental Fig 7B-C, (E) cell viability as the percentage of annexinV and 7AAD negative cells by flow cytometry with or without GGPP (n=5), (F) basal oxygen consumption rate in pmol/minute as assessed by Seahorse technology with or without GGPP (n=8), (G) maximum uncoupler-induced oxygen consumption rate in pmol/minute as assessed by Seahorse technology with or without GGPP (n=8), (H) representative Seahorse tracings presented as oxygen consumption rate (pmol/min) for Supplemental Fig 7F-G, (I) TOMM20 presented as MFI in viable cells (by FVS-R) with or without GGPP (n=4), (J) mitochondrial ROS presented as the ratio to vehicle of MFI of MitoSox in viable cells (by FVS-R) with or without GGPP (n=3), (K) total glutathione presented as uM per 25,000 cells with or without GGPP (n=4), (L) ratio of reduced to oxidized glutathione (GSH/GSSG) assessed in 25,000 cells treated with or without GGPP (n=4), and cell viability as the percentage of annexinV and 7AAD negative cells by flow cytometry with or without (M) GGTI298 (n=4), (N) GGTI2133 (n=3), or (O) FTI277 (n=3). Statistical analysis by Student's T Test. p-values: * = <0.05, ** = <0.01, *** = <0.001. n is the number of replicates.

**Supplemental Table Legends**

**Supplemental Table 1:** (A) Clinical and demographic characteristics of primary AML samples included in the RNA sequencing experiment. Summary of (B) differential gene expression and (C) GSEA in primary *TP53*$^{mut}$ (n=9) *versus* *TP53*$^{WT}$ (n=21) AML patient samples. (D) Additional gene sets included in the GSEA. (E) Clinical and demographic characteristics of the 364 patients evaluated in the retrospective chart review. (F) Summary of cell line characteristics, including focused *TP53* sequencing of the isogenic *TP53*$^{mut}$ and *TP53*$^{WT}$ AML cell line clones.

Supplemental Figure 1:

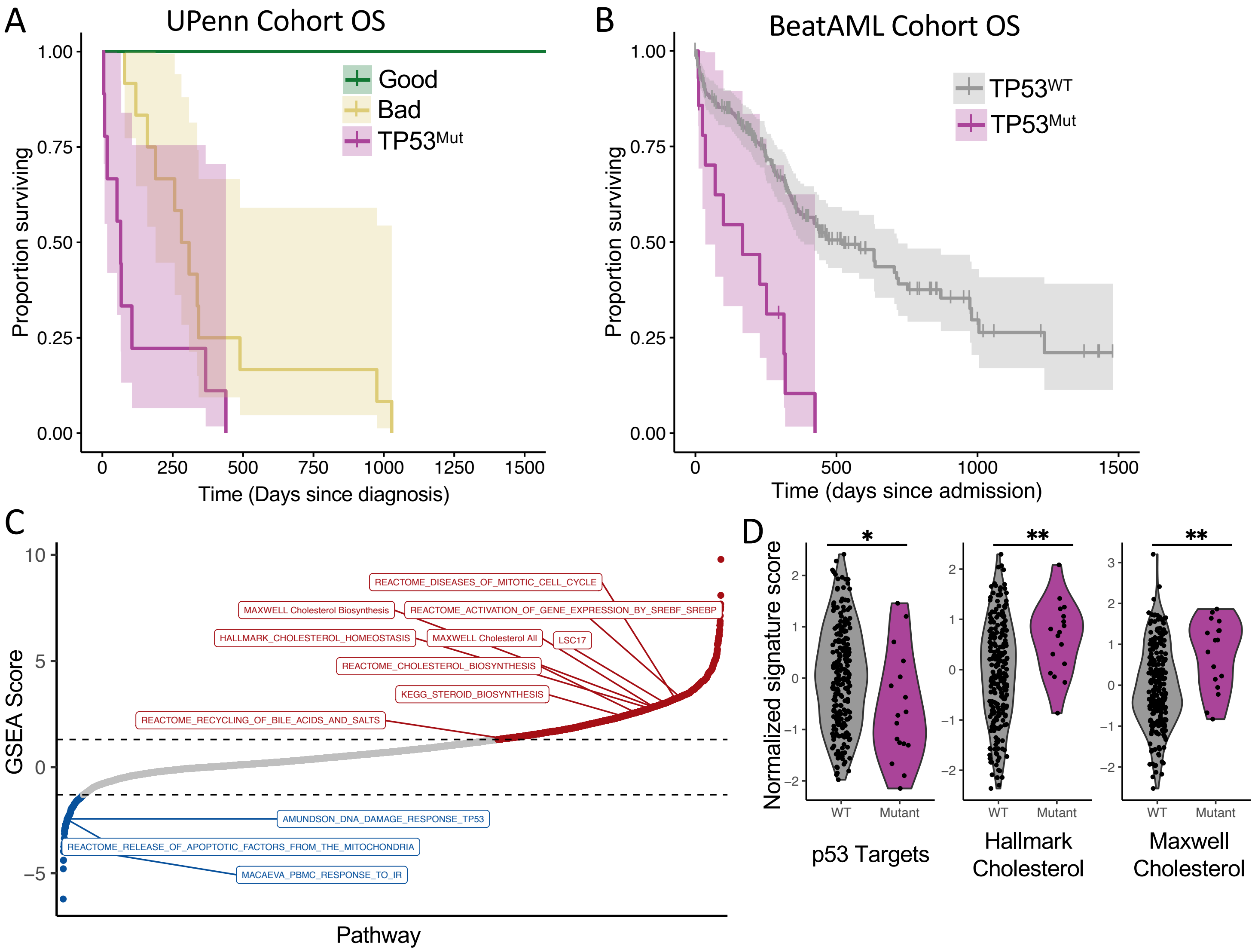

Supplemental Figure 2:

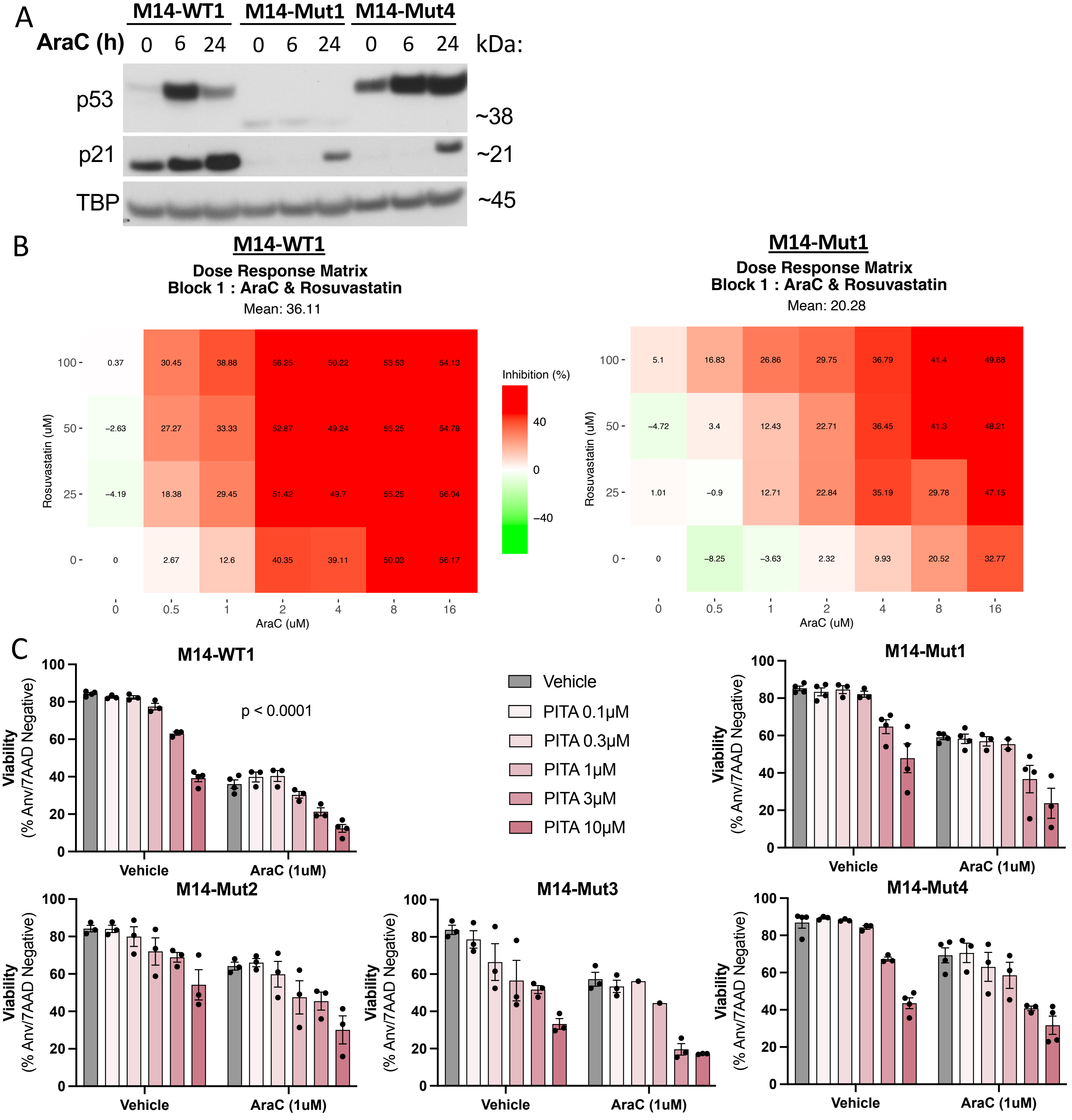

Supplemental Figure 2 Continued:

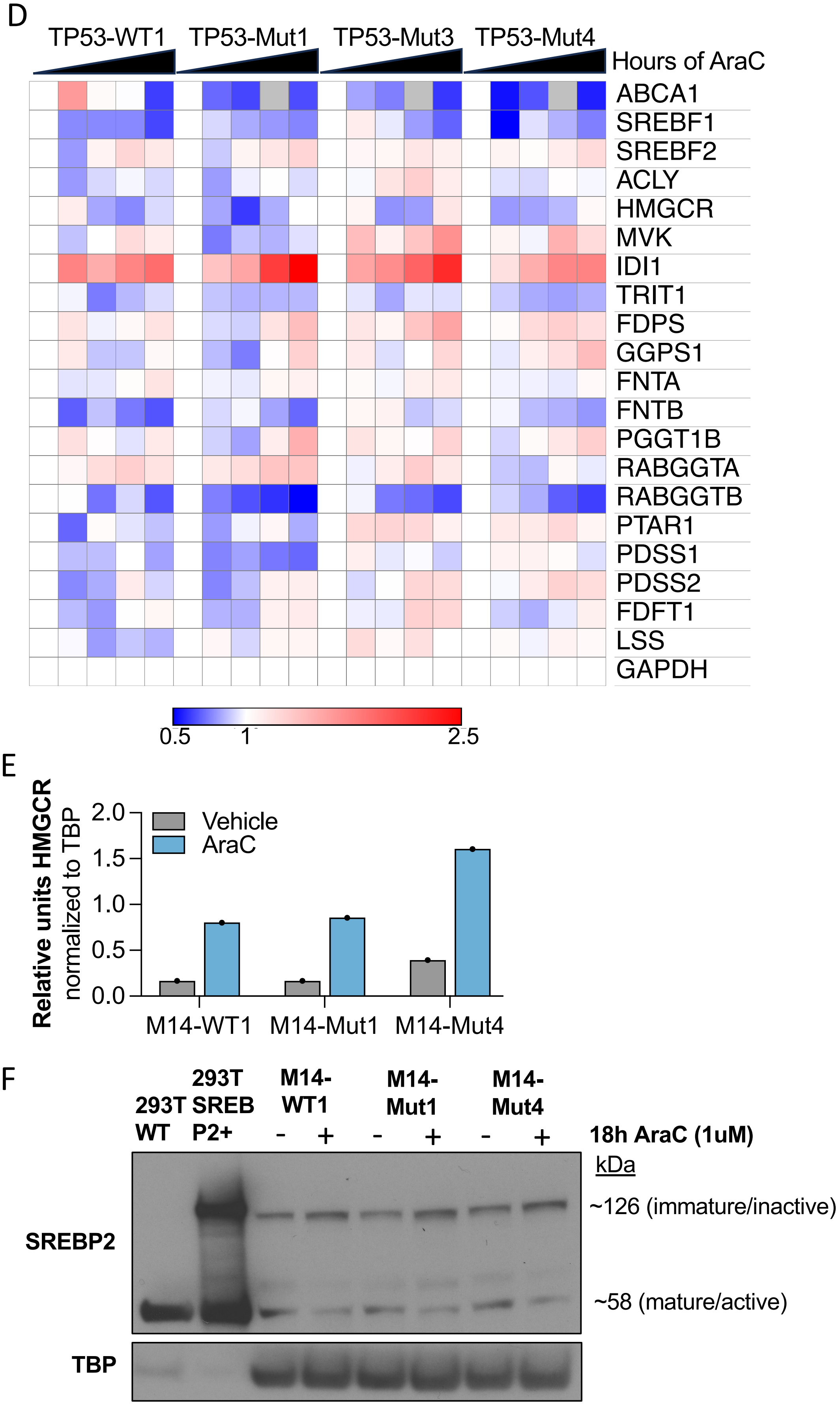

Supplemental Figure 3:

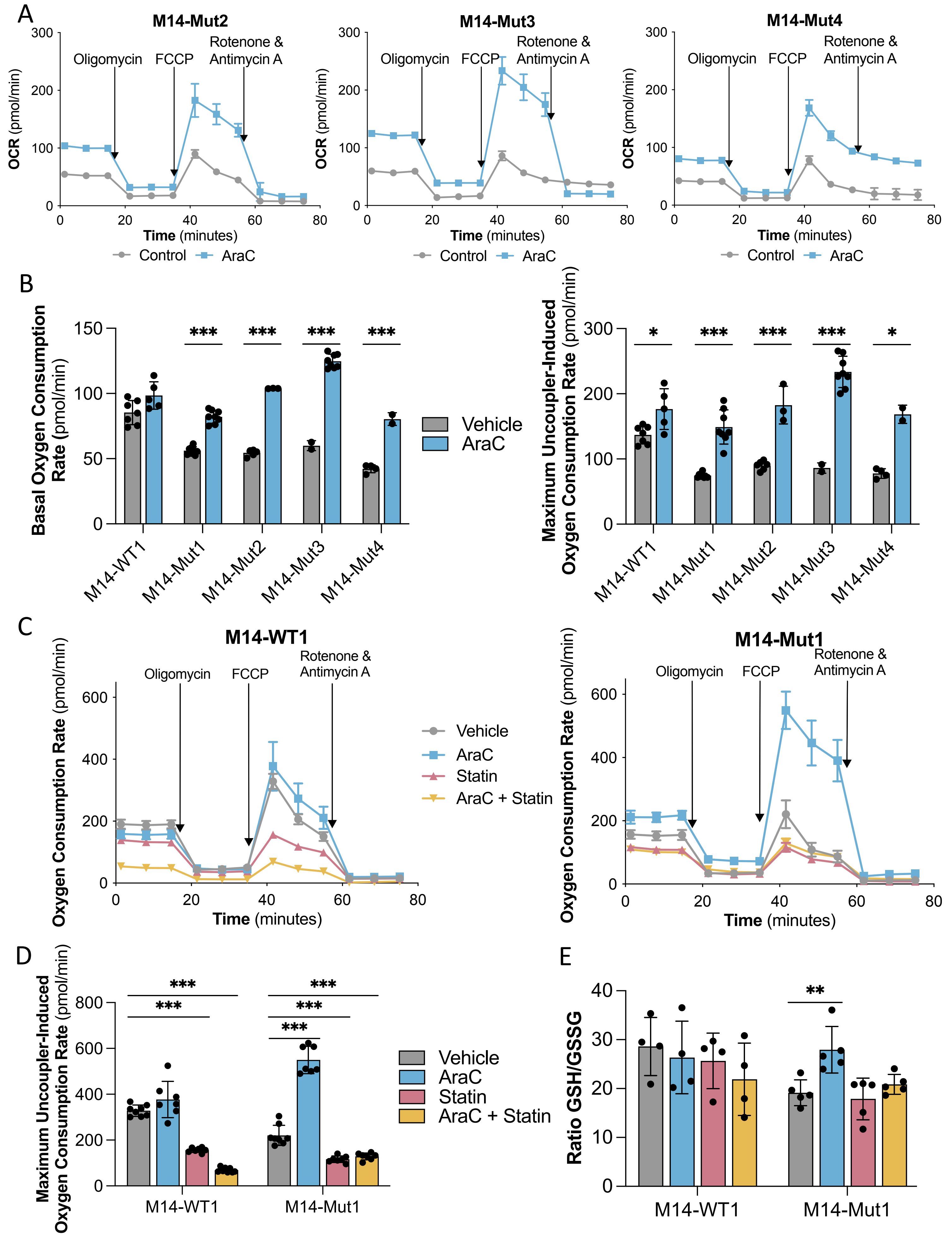

Supplemental Figure 4:

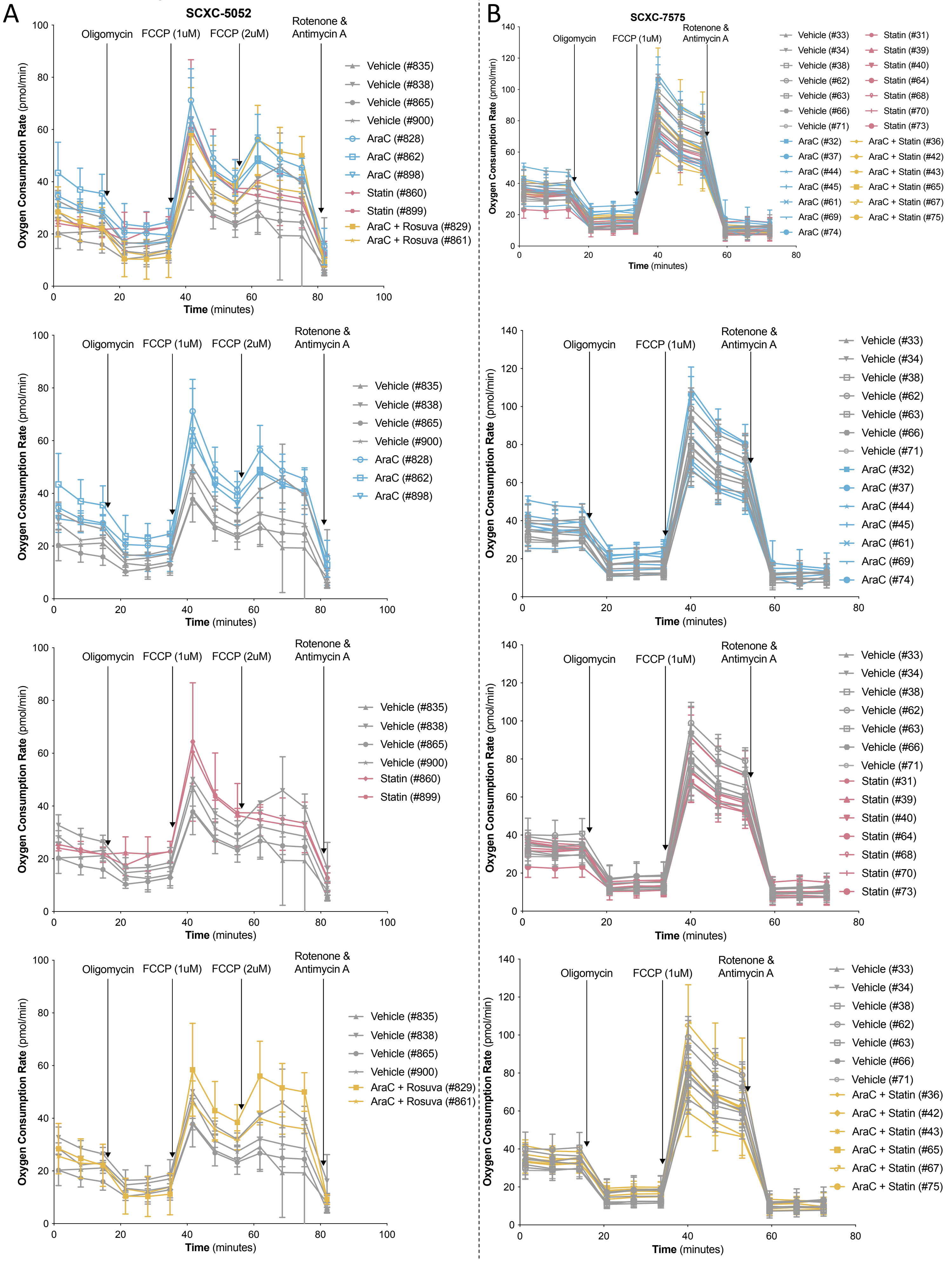

Supplemental Figure 5:

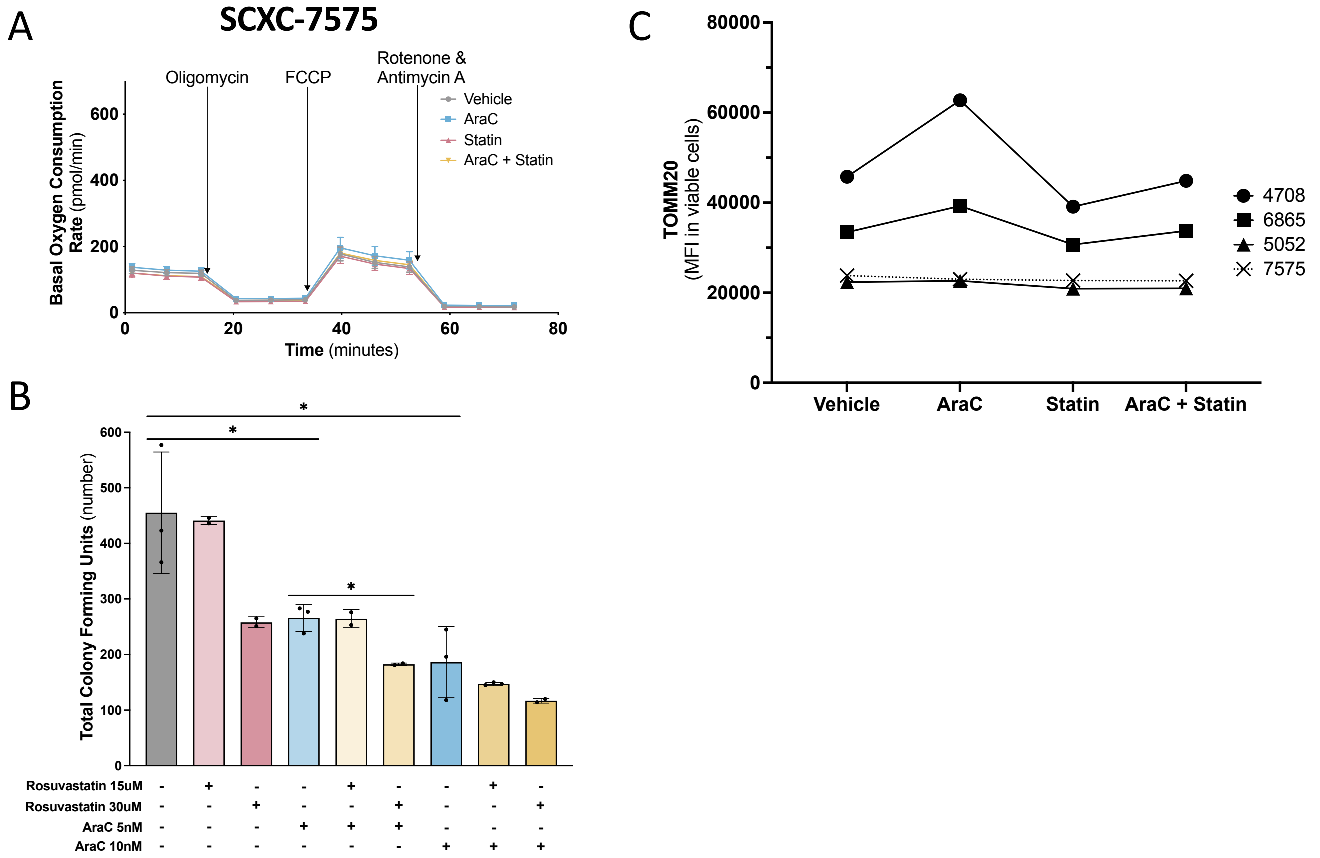

Supplemental Figure 6:

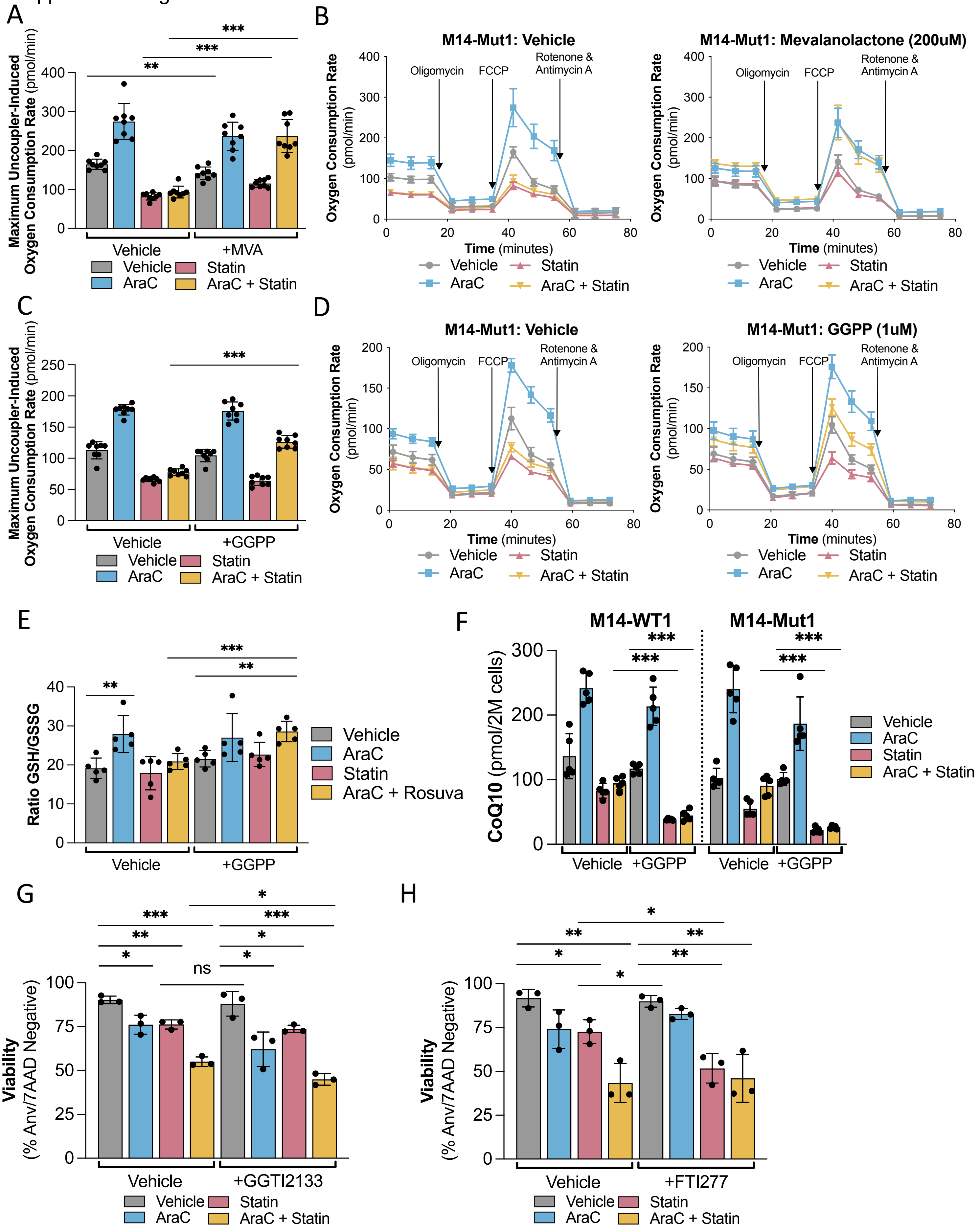

Supplemental Figure 7:

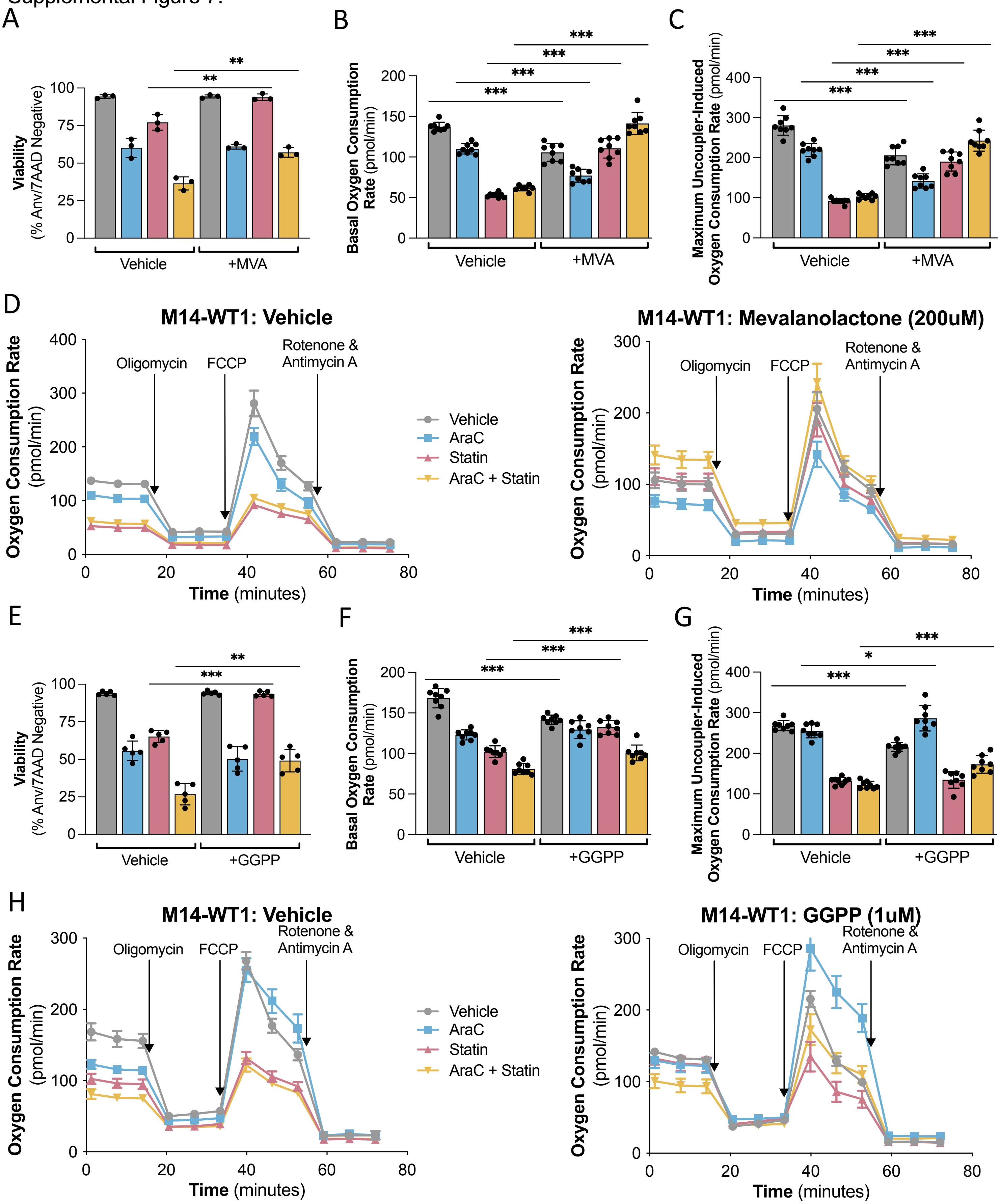

Supplemental Figure 7 Continued:

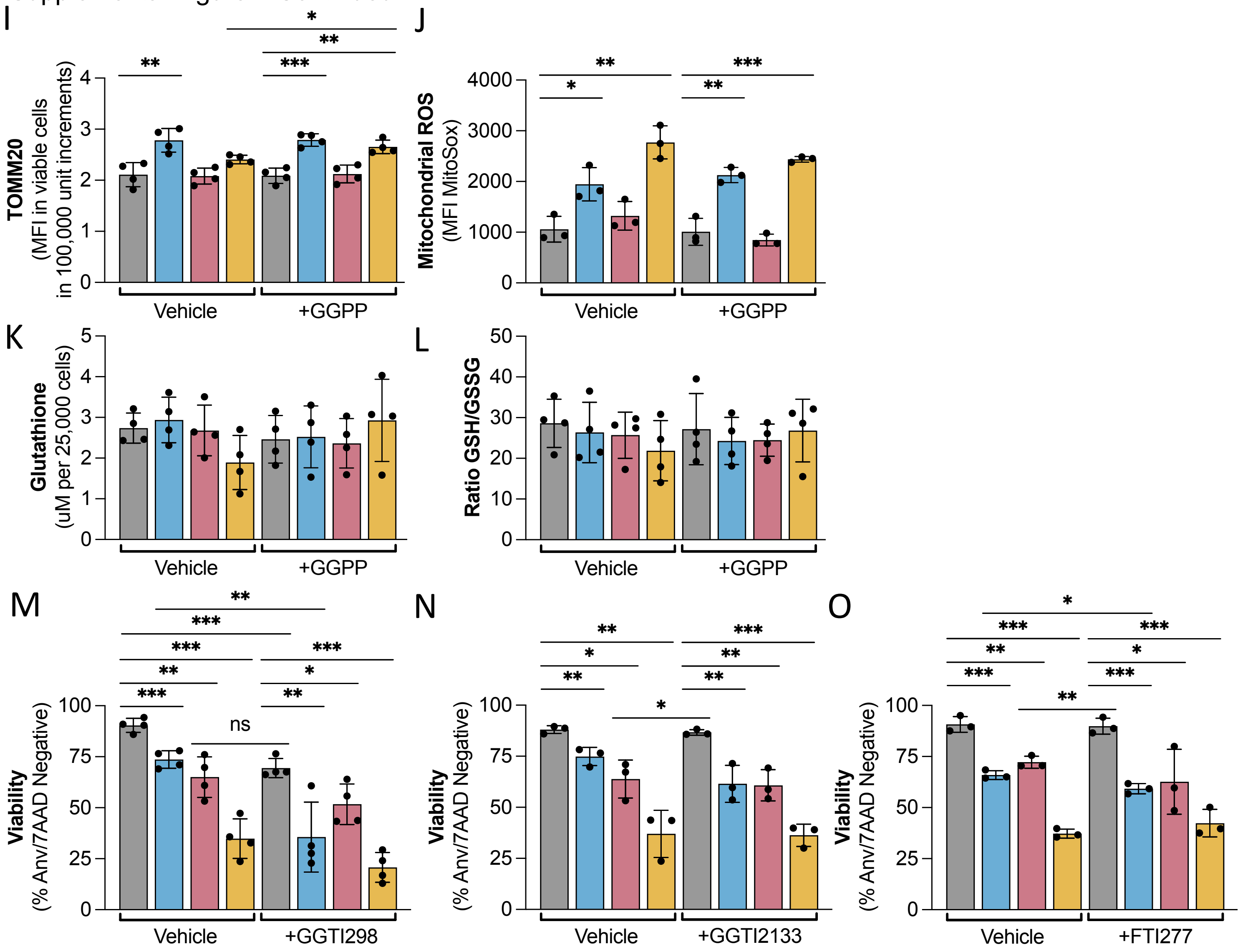
